# CD8 T cells mediate immunosurveillance for neoantigen+ epithelial stem cells in the colon

**DOI:** 10.1101/2025.10.19.683142

**Authors:** Jessica Buck, Nivedita R. Iyer, Eric Fagerberg, Goran Micevic, Jiaming He, Kerri St. Denis, Vyom Shah, Holly Nicole Blackburn, Srividhya Venkatesan, John Attanasio, Brian G Hunt, Alex Dalrymple, Liza Konnikova, Semir Beyaz, Martina Damo, Carmen J. Booth, Kelli A. Connolly, Nikhil S. Joshi

## Abstract

Epithelial cells in the colon accumulate substantial numbers of somatic mutations, some of which can be recognised as neoantigens. The ability of CD8 T cells to survey for neoantigen+ cells in the healthy colon would provide an early detection mechanism to prevent cancer, but it is unclear whether neoantigen-specific CD8 T cells can mediate this process of immunosurveillance without becoming tolerant. To address this question, we used a genetically engineered mouse model to express a neoantigen in the epithelial cells of the adult proximal colon. Induction of neoantigen expression led to rapid elimination of neoantigen+ epithelial cells from the colon in a CD8 T cell-dependent manner. Neoantigen-specific CD8 T cells acquired cytolytic function within the colon tissue under steady-state conditions, which was required for elimination of the neoantigen+ epithelial cells. Despite the elimination of ∼25% of their epithelial cells over a two-day period, the colons looked histologically normal. Immunofluorescence and single-cell transcriptomic analyses revealed that neoantigen-specific CD8+ T cells specifically target neoantigen+ stem cells at the crypt base, which was associated with Ki67 in the crypt wall and abundance of neoantigen-negative stem cells. Infiltrating neoantigen-specific CD8 T cells made IFNg and expressed PD-1, raising the question of why PD-1-dependent suppression did not prevent the acquisition of effector functions by these neoantigen-specific CD8 T cells. Despite an increased signature of interferon-stimulated genes in colonic epithelial cells, PD-L1 expression was surprisingly absent. Moreover, we found that colonic epithelial stem cells also did not express PD-L1 under conditions of chronic inflammation, such as ulcerative colitis, immune checkpoint-induced colitis, and ageing, or when directly stimulated with IFN-γ *in vitro*. Analyses of the PD-L1 gene promoter across humans and mice showed hypermethylation at sites associated with PD-L1 repression in cancer. Thus, our data support a model in which the acquisition of neoantigens by colonic epithelial cells triggers CD8 T cell-mediated immunosurveillance. This results in the elimination of PD-L1-negative neoantigen+ stem cells by effector CD8 T cells and simultaneous repair of the colon by neoantigen-negative epithelial cells to prevent immunopathology.

## Introduction

Cells in barrier tissues, including the skin, lung, and intestine, accumulate somatic mutations at a high rate due to exposure to environmental mutagens, which presents an increased risk for transformation ^1, 2, 3, 4^. Some mutations can create neoantigens that are presented on MHC class I, providing a means for the immune system to distinguish and potentially eliminate the mutated cells ^5^. In the colon, it is also necessary to establish and retain immunologic tolerance to non-harmful sources of neoantigens, like those from commensal bacteria and food ^6, 7, 8^. Inability to establish peripheral tolerance results in intestinal inflammatory conditions, such as ulcerative colitis. So, the prevailing view is that the colon tissue microenvironment is dominated by immunosuppression ^9, 10^. The intimate and long-term interactions between the intestinal epithelium, luminal bacteria, and environmental mutagens result in a healthy colon epithelium with high numbers of neoantigens ^11, 12, 13, 14, 15, 16^. CD8+ T cells are uniquely able to both recognise neoantigens and infiltrate peripheral tissues to survey large numbers of parenchymal cells and thus are ideal candidates for mediating the immunosurveillance necessary for maintaining homeostasis in the colon ^17, 18, 19, 20^. However, it remains unclear whether or how neoantigen-specific CD8 T cells that retain functional potential surveil healthy colon tissue at steady state.

Before CD8 T cells can expand and survey, they must be primed in lymphoid tissues via interactions with neoantigen-presenting dendritic cells (DCs)^21^. In the absence of overt inflammation, DC-CD8 T cell interactions in lymphoid tissues yield peripheral tolerance^22, 23^. Mechanistically, these CD8 T cells are rendered non-functional via anergy-induction in the lymphoid tissue, or die due to apoptosis, preventing these cells from entering peripheral tissues and mediating immunopathology ^24^. Regulatory T cells (Tregs) also play a key role in this process by suppressing self-reactive CD8 T cell activation. Yet, tolerance in this manner would prevent CD8 T cells from mediating immunosurveillance in the tissue, highlighting a key gap in our understanding of how immunosurveillance is achieved while self-tolerance is maintained.

Studies of CD8 T cell-immunosurveillance or self-tolerance have traditionally been performed in animal models expressing model antigens. However, most models have promiscuous expression of the model antigen in the thymus^25^. Leaky antigen expression results in the deletion of endogenous antigen-specific CD4 and CD8 T cells and can also lead to the generation of large numbers of endogenous antigen-specific Tregs.^26, 27^ This highlights the potential confounding impact of central tolerance mechanisms on prior studies of peripheral tolerance. To circumvent these complications, we previously established the iNversion INducible Joined neoAntigen (NINJA) model, where splicing and DNA inversion result in neoantigens that do not exist in the thymus or peripheral tissues until induced^28^. These neoantigens are derived from lymphocytic choriomeningitis virus (LCMV) and are contained within a loop of green fluorescent protein (GFP). In the OFF state, all cells express just the N-terminal portion of GFP, but are not fluorescent. After induction, recombination events result in cells that express full-length GFP that contain the neoantigens. Consequently, NINJA mice are tolerant to GFP but feature a full repertoire of neoantigen-specific naïve CD8 and CD4 T cells.

Induction of NINJA is highly regulated, requiring expression of Cre recombinase and Doxycycline and Tamoxifen (Dox/Tam, D/T) treatment. Dox/Tam administration induces neoantigen expression in Cre-expressing cell types, enabling us to study the impact on endogenous neoantigen-specific CD8 and CD4 T cells. Using this approach, we previously identified a mechanism by which PD-1 blockade disrupts skin-specific CD8 T cell tolerance. PD-1 blockade resulted in T cell killing of NINJA+ keratinocytes and caused immunopathology, unexpectedly diverging from classical definitions of T cell tolerance.^29^ Thus, by leveraging the strengths of the NINJA system, it is possible to refine studies of tolerance and to reveal previously unappreciated T cell immunosurveillance mechanisms.

Here, to gain mechanistic insights into T cell immunosurveillance in the colon under homeostatic conditions, we used NINJA to induce neoantigen expression under the control of a promoter expressed in proximal colon epithelium. This allowed us to study how endogenous, neoantigen-specific CD8 T cells mediate immunosurveillance and ultimately eliminate neoantigen-expressing epithelial cells without causing immunopathology. These studies offer insights into the mechanisms that enable the efficient clearance of neoantigen+ cells, providing a potential explanation for why immune-related adverse events (irAEs) in the colon are far less common after PD-1/PD-L1 blockade than in other tissues.

## Results

### Neoantigen-expressing colon epithelial cells are eliminated in a CD8 T cell-dependent manner

To study T cell tolerance in the colon, we established a system to inducibly express a neoantigen in colon epithelial cells. We first needed to identify a Cre recombinase that provided inducible expression in the colon. CDX2-CreERT2 is known to be most active in the proximal colon epithelium and is inducible by treatment with Tamoxifen (Tam, T)^30^. To confirm inducibility, we bred CDX2-CreERT2 to Ai14 mice (CAi14 mice), which contain a *loxP*-STOP-l*oxP*-tdTomato cassette. We treated mice with Tam (200 μL, 20 mg/mL provided by oral gavage) and 15 days later, we sacrificed animals, removed their colon, and analysed the gross expression of tomato using an Xite Portable Fluorescence Flashlight (**Fig. 1a**). This showed that Cre activity was primarily restricted to the proximal colon and caecum, with decreasing activity towards the mid colon and little-to-no activity in the distal colon. We further analysed tomato expression by immunofluorescence on PFA-fixed and frozen colon sections, where the colon had been rolled from distal (inside) to proximal (outside) (**Supplementary Fig. 1a**). Note, in the mouse, the proximal colon has large transverse folds which are not present distally. We were able to identify tomato+ crypts (where the entire crypt from the base to the top was comprised mostly of tomato+ cells), and the majority of the crypts in the proximal colon were tomato+. In the mid-colon, the presence of tomato+ crypts became patchier, with “neighbourhoods” of positive crypts surrounded by larger regions of tomato-negative crypts. In the distal colon, there were few, if any, tomato+ crypts, in line with the lack of tomato seen grossly via the Xite fluorescence flashlight. We confirmed that tomato expression in the proximal colon was stable over 12 days (**Supplementary Fig. 1b**). Given epithelial cell turnover is three to five days in the mouse colon, this suggests we had permanently labelled a subset of epithelial stem cells in the tissue, consistent with prior studies on CDX2-CreERT2^30^.

**Fig. 1:**
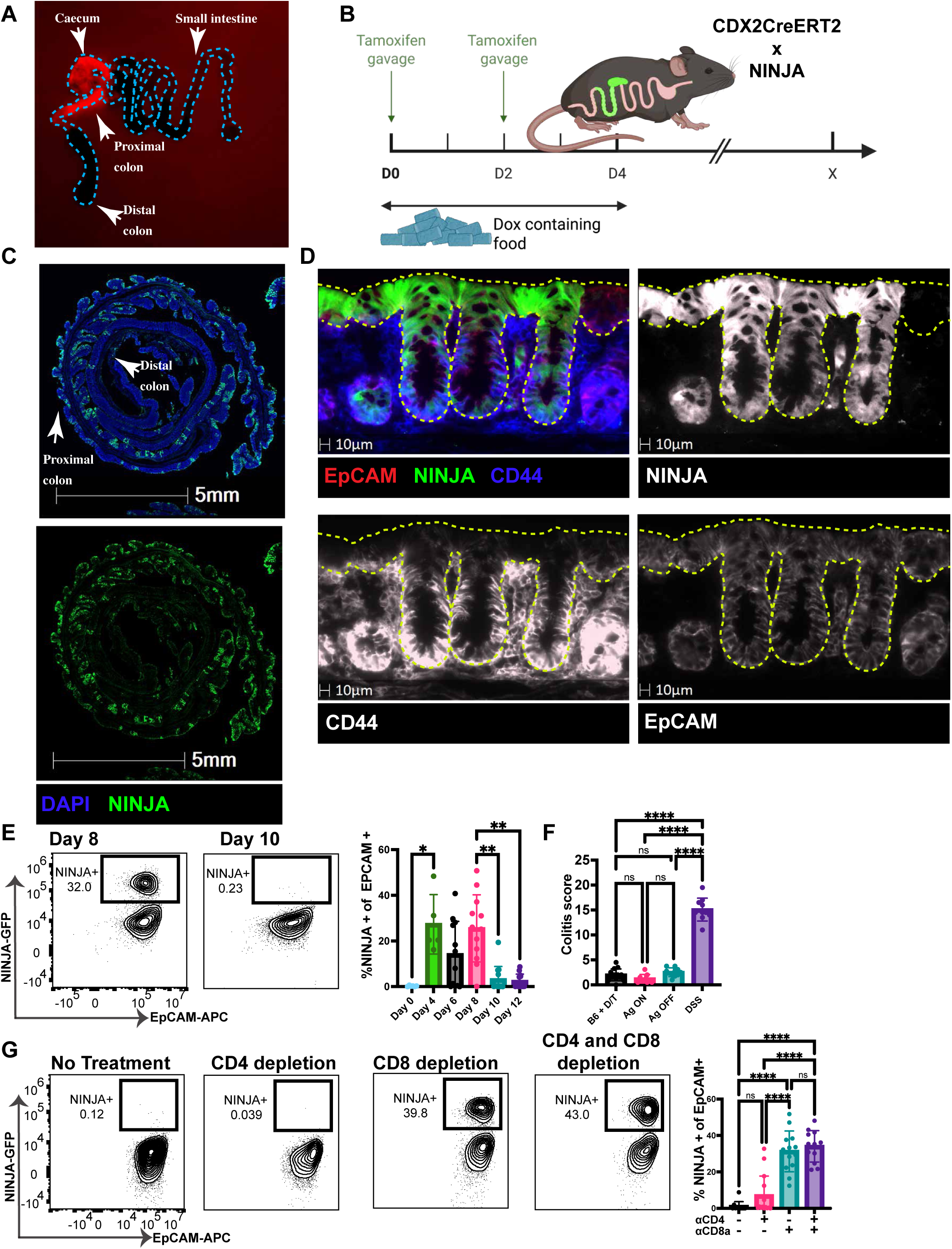
Antigen-expressing epithelial cells are lost in a T-cell-dependent manner. **a**,Photograph of intestines 6 weeks post gavage using green Xite fluorescent flashlight system, n = 6 across two experimental repeats. The blue dashed line outlines the intestine. **b**, Antigen induction and experimental schedule schematic. **c**, 20x CellDIVE immunofluorescence image of Swiss-rolled colon proximal (outside) to distal (inside) on Day 4, n = 7 across 2 experimental repeats. blue = DAPI, green = NINJA. **d,** 20x CellDIVE immunofluorescence image on Day 6, n = 8 across 2 experimental repeats. Dotted yellow line denotes epithelium. Pink = EpCAM, green = NINJA, blue = CD44. **e**, Ninja expression time course. Frequencies of NINJA-expressing proximal colon epithelial cells. Data are mean ± s.d. n = 6 for day 0, n = 5 for day 4, n = 12 for days 6 and 8, n = 13 for days 10 and 12. n is reported as the total across three experimental replicates except for day 4, which was included in two experimental replicates. **f**, Colitis scores from H&E (Hematoxylin and Eosin)- and PAS (Periodic Acid Shiff)-stained sections of proximal mouse colon from DSS colitis control mice (n = 10) with loss of crypts, inflammation, and edema compared to unremarkable proximal colon from mice with antigen on (n = 9), antigen off (n = 10), and C57BL/6 treated with dox/tam (n = 12). n is reported as the total across three experimental replicates, except for the DSS group, which is two experimental replicates **g**, Flow cytometry and quantification of NINJA+ epithelial cells day 12 post antigen induction from no treatment (n = 11), CD4+ T cell depletion (n = 15), CD8+ T cells depletion (n = 13), and both CD4+ and CD8+ T cell depletion (n = 12). n is reported as the total across three experimental replicates. Kruskal-Wallis test with Dunn’s multiple comparison test was used to determine statistical significance of the NINJA time course. One-way ANOVA with Tukey’s multiple comparison test was performed to determine the statistical significance of the colitis scores and the T cell depletions. For all statistical tests, *P < 0.05; **P < 0.01; ***P < 0.001; ****P < 0.0001

Having established the pattern of CDX2-CreER expression in the colon, we next bred mice containing CDX2-CreERT2 to NINJA x Cag-rtTA3, to generate F1 mice containing all three alleles (referred to as CNC mice) (**Fig. 1b**). In NINJA, the LCMV-derived peptides GP33-43 and GP60-80 are contained within a loop of GFP. In the NINJA off configuration, which is the baseline for every cell in the mouse, a truncated GFP product is made that is not fluorescent and does not contain the neoantigens^28^(**Supplementary fig. 1c**). In the presence of Cre recombinase, doxycycline (Dox, D), and tamoxifen, a series of recombination events results in full-length GFP containing the neoantigens. Thus, GFP expression marks neoantigen+ cells^28^. For simplicity, we will refer to the neoantigen-containing GFP+ colon epithelial cells in CNC mice as “NINJA+” cells, while GFP-cells in the OFF state will be referred to as “NINJA-negative” cells.

Treating CNC mice with D/T (dox from days 0-4 provided in food and Tam treatment on days 0 and 2 provided by gavage) resulted in NINJA+ epithelium, mainly in the proximal colon (**Fig. 1b-c**). By day 6, NINJA was expressed throughout individual crypts, with notable expression at the crypt base (defined by high CD44 expression) (**Fig. 1d**). This pattern of expression is consistent with stem cell expression of NINJA, which we subsequently confirmed via scRNA-Seq (see below), and the known CDX2-Cre-ERT2 activity (**Fig. 1a, Supplementary fig. 1a-b**)^30^.

To quantify the fraction of NINJA+ epithelial cells in the proximal colon, we isolated, digested, and prepared proximal colon tissue to enrich for epithelial cells, then analysed EpCAM+ epithelial cells for GFP expression by flow cytometry (FACS; **Fig. 1c, Supplementary fig. 1d**). At day 8, approximately 25% of the proximal colon epithelial cells were NINJA+, but strikingly, by day 10 only about 3% were NINJA+. We confirmed the loss of GFP expression using time-course analyses between day 4 and 12, which demonstrated the presence of NINJA+ epithelial cells between days 4-8 and then absence on days 10-12 (**Fig. 1e**). Notably, asynchronous induction due to feeding and gavage over a period of several days led to some variability between mice with respect to kinetics of neoantigen-induction. Nonetheless, despite a substantial loss of epithelial cells, the CNC mice showed no overt pathology (**Fig. 1f and Supplemental fig. 1e**).

To test if the loss of NINJA was due to a T cell response, we depleted either CD4, CD8, or both CD4 and CD8 T cells with anti-CD4 or anti-CD8a antibodies, starting with treatment on days −14 prior to NINJA induction. We confirmed, via FACS, that the antibody treatment was effective at depleting either CD8 or CD4 T cells by staining for T cells using anti-CD8b or anti-CD4 (clone RM4-5) clones that are not blocked by the depletion antibodies (**Supplemental fig. 1f**). Depletion of CD8 T cells and both CD4 and CD8 T cells led to significant increases in the fraction of NINJA+ epithelial cells on day 12 post induction (**Fig. 1g**). By contrast, depletion of CD4 T cells alone did not significantly increase the fraction of NINJA+ epithelial cells. However, a handful of mice did retain some NINJA+ epithelial cells (partial rescue of NINJA expression in 3/15 and full rescue in 3/15), which contrasted with non-depleted mice (partial NINJA expression in 1/11) (**Fig. 1g**). Thus, NINJA+ colon epithelial cells are cleared between days 8 and 10 post neoantigen induction in a CD8+ T cell-dependent manner.

### GP33-specific CD8+ T cells are cytolytic effectors

Given that CD8 T cells were required for loss of the NINJA+ epithelial cells, we next assessed the infiltration, phenotypes, and functions of neoantigen-specific CD8 T cells. We focused our analyses on day 8, as this was the last day NINJA+ epithelial cells were routinely observed. We performed FACS on isolated lymphocytes from the proximal colon lamina propria (LP), intraepithelial layer (IEL), and mesenteric lymph nodes (MLNs). Neoantigen-specific CD8 T cells were detected using GP33-peptide-loaded Db-MHC class I tetramers (**Supplemental fig. 2a**). There was a significant increase in the fraction of GP33-specific CD8+ T cells in the MLNs, LP, and IEL when NINJA was induced (**Fig. 2a**). Analysis of phenotypic markers on the GP33-specific CD8 T cells expressed activation markers CD44 and PD-1 in all tissue sites. In the IEL, GP33-specific CD8 T cells showed higher expression of the activation and tissue-residence markers CD69 and CD103 compared to the cells in the MLN (**Fig. 2b-d, and Supplemental fig. 2b**). GP33-specific CD8 T cells also had a progressive increase in Granzymes A and B expression between the MLN, LP, and IEL, with nearly all cells in the IEL expressing Granzyme A (**Fig. 2e-f**), a marker of terminal effector CD8 T cells ^31, 32^. Notably, GP33-specific CD8 T cells in all sites were perforin+, suggesting a high capacity for cytolysis, with the highest frequency of perforin expression in the colon tissue. This suggested that GP33-specific CD8+ T cells in the IEL are cytolytic effector T cells that acquired effector function in the colon tissue.

**Fig. 2:**
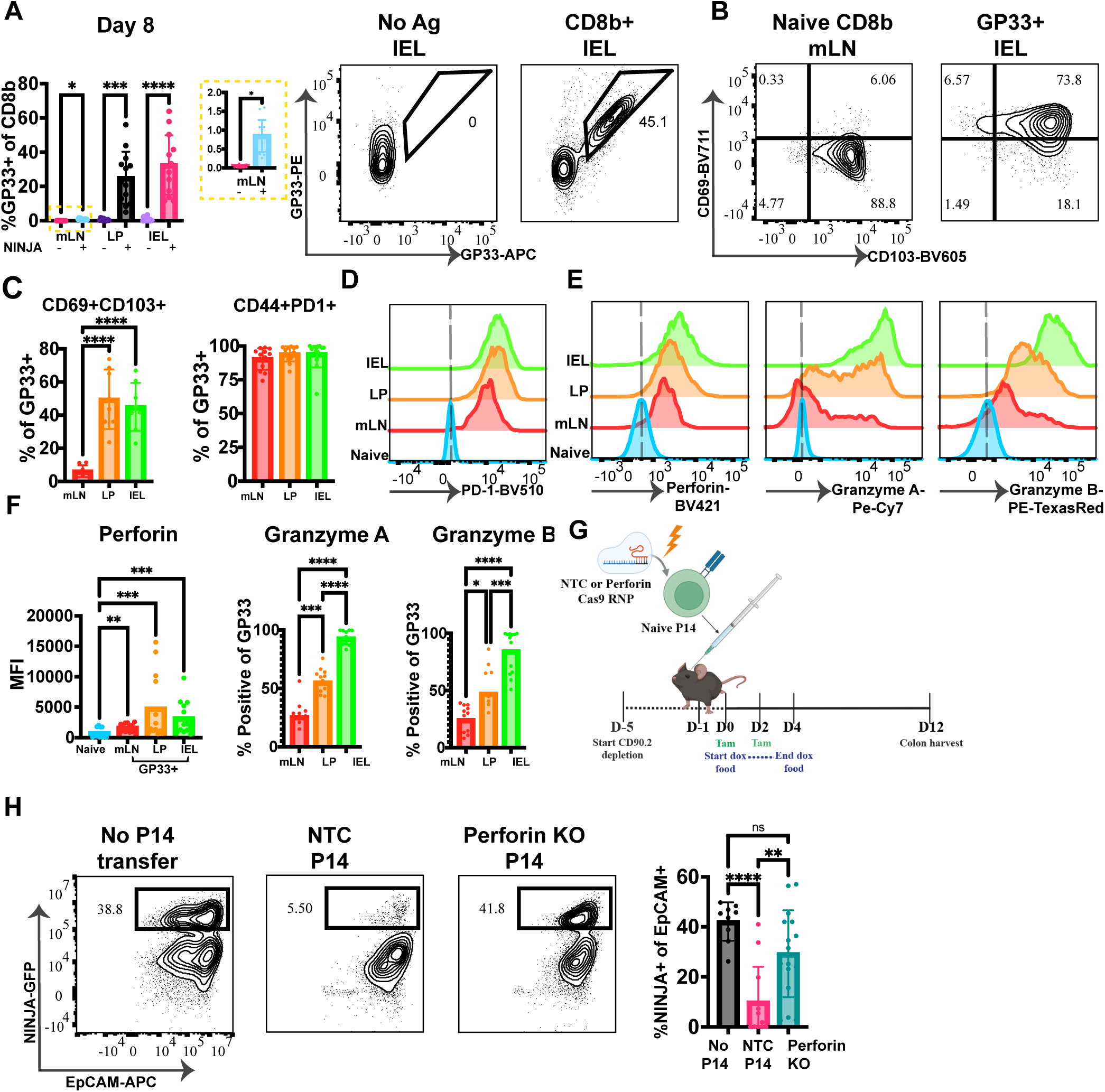
NINJA-specific T cells are cytolytic and clear NINJA+ epithelial cells via Perforin. **a-f**, Flow cytometry phenotyping of GP33-specific CD8b+ T cells, day 8 post-NINJA induction. n = 13 for Ninja on and n = 12 for NINJA off, across 3 experimental repeats. **a,** Frequencies of NINJA-specific CD8+ T cells within the mLN, LP, and IEL on day 8 and antigen-specific gating strategy. **b**, Representative gating strategy for CD69 and CD103+ NINJA-specific CD8+ T cells. **c**, Frequency of CD69 +and CD103+ as well as CD44+ and PD1+ NINJA-specific CD8+ T cells, with a representative PD-1 histogram **d**, representative PD-1 histogram. **e**, representative histograms of Perforin, Granzyme B, and Granzyme A. **f**, MFI quantification of Perforin expression and percent of GP33+ CD8b+ T cells expressing granzyme B and granzyme A. **g,** Graphical depiction of perforin KO experimental timeline. **h,** Representative flow cytometry plots depicting %NINJA+ epithelial cells with quantification of NINJA+ epithelial cells. Kruskal-Wallis test with Dunn’s multiple comparisons was used to determine statistical significance of the phenotyping data. Ordinary one-way ANOVA test with Tukey’s multiple comparison test was used to determine statistical significance of the perforin KO experiment. For all statical tests, *P < 0.05; **P < 0.01; ***P < 0.001; ****P < 0.0001.

To test if colon epithelial cells were killed by cytolytic, GP33-specific CD8 T cells, we next depleted all endogenous T cells from CNC mice using antibodies against Thy1.2 (CD90.2) and then reconstituted with an adoptive transfer of naïve Thy1.1+ GP33-specific T cell receptor (TCR)-transgenic P14 CD8 T cells, which were insensitive to the anti-Thy1.2. This system allowed us (1) to establish that GP33-specific CD8 T cells were sufficient to eliminate the neoantigen+ epithelial cells, and (2) to knock out genes required for cytolytic function in the transferred naïve P14 CD8 T cells to determine the necessary molecules for elimination of neoantigen+ epithelial cells. As CD8 T cell cytolytic function via Granzymes is dependent on perforin, we electroporated naïve P14 T cells with Cas9 protein complexed to sgRNAs specific for perforin or non-target controls (NTCs; controls)(**Fig. 2g**). Critically, this protocol does not activate the CD8 T cells prior to adoptive transfer^33^. As expected, depletion of all T cells with anti-Thy1.2 resulted in the maintenance of NINJA+ EpCAM+ epithelial cells at day 12 post-induction, which were eliminated by the transfer of unmodified (NTC) Thy1.1+ P14 CD8 T cells (**Fig. 2h**). Thus, the adoptively transferred naïve GP33-specific CD8 T cells were sufficient to eliminate the neoantigen+ colonic epithelial cells. Importantly, perforin knockout in the transferred GP33-specific CD8 T cells abrogated their ability to kill the neoantigen+ colonic epithelial cells, demonstrating that the cytolytic capacity of neoantigen-specific CD8 T cells was dependent on the expression of perforin (**Fig. 2h**).

### Colon-reactive CD8 T cells are primed in colic MLNs

An alternative explanation to the above hypothesis that T cells gain cytolytic function in the colon could be that GP33-specific CD8 T cells are primed outside of the MLN. To confirm that the proximal colon MLN was the site of T cell priming, we assessed sites of early TCR signalling in the MLN shortly after antigen induction. The colon and small intestine (SI) drain into distinct LNs within the mesenteric lymph node chain, allowing us to compare priming within the cMLNs (colon draining MLN) to the sMLNs (small intestine MLNs), which are neighbours ^34, 35^. To assess TCR signalling, we made use of Thy1.1+ GP33-specific P14 TCR transgenic mice bred to *Nr4a1*-GFP (*Nr4a1* encodes the Nur77 protein) transgenic reporter mice (henceforth called Nur77-reporter P14 mice). Thy1.1+ GP33-specific CD8 T cell-containing splenocytes were isolated from Nur77-reporter P14 mice, labelled with cell trace violet (CTV) dye to assess proliferation, and adoptively transferred into Thy1.2+ CNC mice. On days 2 and 3 after induction, we assessed proliferation and TCR signalling on the transferred P14 T cells in the cMLNs, sMLNs, and negative control inguinal LNs (iLNs; **Supplementary fig. 3a,b**). On day 2, Nur77 was most highly expressed by P14 T cells in the cMLNs (60.7%; SD 10.1). In sMLNs, 25.8% (SD 7) of P14s were Nur77+ and 4.3% (SD 0.8) were Nur77+ in the iLNs. Moreover, by day 3, 40.1% (SD 6.5) of the P14 T cells had divided at least 5 times in the cMLNs compared to 16.6% (SD 3.4) and 4.7% (SD 1.8) in the sMLNs and iLNs, respectively. Altogether, these data demonstrate that priming primarily occurred in the cMLNs, with cells subsequently migrating to other sites.

To confirm that antigen-presenting cells (APCs) were draining from the colon to the cMLNs, we again leveraged Cai14 mice to induce tomato expression in colon epithelial cells and track accumulation of tomato+ macrophages (MF) and dendritic cells (DCs) in different sites (**Supplementary fig 3c,d**). Cre expression determines which epithelial cells express NINJA, so in Cai14 mice, tomato is expressed on the same cells. Tomato is a pH-stable protein that we and others have previously used to track uptake of cellular material^36, 37^. On day 6 after induction, in the tissue (LP), we noted that Xcr1+ and Xcr1-DCs, as well as F4/80+ Mφ, had taken up tomato. However, in the MLNs, the only tomato+ cells were Xcr1+ DCs (**Supplementary fig 3e**). Critically, amongst the different MLNs, tomato was only found within XCR1+ dendritic cells in the cMLN, and not the sMLN (**Supplementary fig 3f**). Together, these data strongly suggest that GP33-specific CD8+ T cells were primed in the cMLN by colon-draining Xcr1+ DCs and then trafficked to the colon, where they acquired full effector functions in the LP and IEL.

### GP33-specific CD8+ T cells infiltrate the crypt base

To understand the interactions between infiltrating CD8 T cells and NINJA+ epithelial cells in the colon, we performed multiplex cyclic immunofluorescence on tissues on days 0-10 post-induction. By day 6, within NINJA expressing crypts, we observed that the infiltrating CD8b+CD3+PD-1+ T cells were all PD-1+, which was consistent with our FACS analysis of the CD8+ T cells at this time point. We also noted that T cells were infiltrating into the epithelium, specifically in NINJA-expressing crypts (**Fig. 3a**). Moreover, a larger fraction of the infiltrating CD8b+ T cells appeared to be enriched at the base of the crypt vs. the top, suggesting preferential localisation (**Fig. 3b**).

**Fig. 3:**
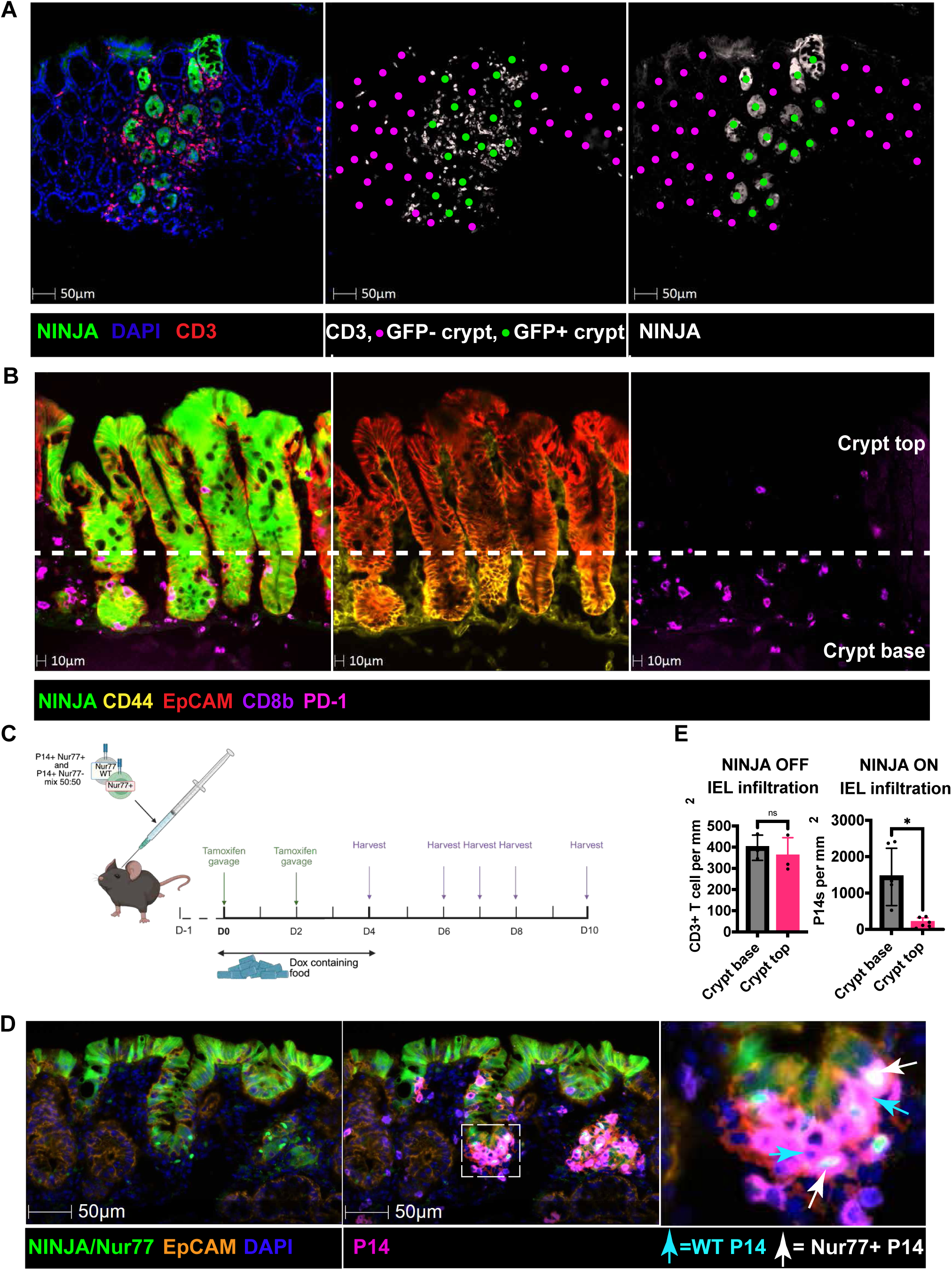
NINJA-specific T cells infiltrate the crypt base. **a**, 20x CellDIVE immunofluorescence image demonstrating endogenous T cell infiltration of NINJA+ crypts. Image is representative of 7 mice, from days 6-8. Green = NINJA, DAPI = blue, and red = CD3. Purple and green dots were manually placed in the centre of NINJA- and NINJA+ crypts, respectively. **b**, 20x CellDIVE immunofluorescence image depicting endogenous CD8b+ PD1+ T cells infiltrating NINJA+ crypt bases. The crypt base is defined by the co-expression of CD44 (yellow) and EpCAM (red), appearing orange. The white dashed line separates the crypt base and top. Green = NINJA, purple = CD8b, pink = PD-1. Image is representative of 7 mice, from days 6-8. **c**, Experimental timeline of Nur77 reporter P14+ T cell transfer. 5 x 10^5^ splenocytes of a 50:50 mix from Nur77 WT and Nur77 reporter mice were transferred 1 day prior to antigen induction. **d**, Representative 20x CellDIVE immunofluorescence image showing P14+ T cell infiltration at the crypt base. Green = NINJA and Nur77. Orange = EpCAM. Blue = DAPI, pink = P14+ T cells (a combination of Thy1.1 (red) and CD8b (purple). The cyan arrows point to Nur77 WT P14+ T cells, and the white arrows point to Nur77 reporter P14+ T cells that are also expressing Nur77. **e**, Quantification of Fig 3d. HaloAI infiltration analysis quantifying the number of endogenous T cells or P14s within the crypt base vs top per mm2 of tissue, before and after NINJA induction, n = 3 for before and n = 6 for after, consisting of mice from days 6-8 with NINJA positivity. Kruskal-Wallis analysis with Dunn’s multiple comparison test was used to determine the significance of the T cell infiltration. *P < 0.05; **P < 0.01; ***P < 0.001; ****P < 0.0001.

To detect if GP33-specific CD8 T cells were seeing antigen based on encounter with NINJA+ epithelial cells, we analysed mice with adoptively transferred, congenically marked naive Nur77 reporter-expressing P14 CD8 T cells. Because the reporter and NINJA were both GFP+, we included congenically marked WT naïve P14 CD8 T cells as controls. This provided confidence that the GFP signal was from the Nur77-GFP reporter in the T cell and not NINJA, because WT T cells should always be GFP-negative. Furthermore, the fluorescence signal from the Nur77 reporter is brighter than NINJA. The mixture of P14 CD8 T cells for adoptive transfer was a 50:50 mix of splenocytes containing Thy1.1+ CD45.2+ Nur77-GFP reporter naïve P14 CD8 T cells and Thy1.1+ CD45.1+ WT (no reporter) naïve P14 CD8 T cells. We transferred 5 × 10^5^ total splenocytes into Thy1.2+ CD45.2+ CNC mice and induced NINJA expression. On days 6 to 8, we analysed the colon tissues by immunofluorescence. Transferred Thy1+ CD8b+ P14 T cells were often located near the base of the NINJA+ crypts (**Fig. 3d**). Many, but not all, Thy1.1+ CD45.2+ cells expressed GFP, suggesting they were receiving TCR signals in the tissue.

As we often observed the neoantigen-specific CD8 T cells were near the base of crypts, we next used HaloAI to quantify the relative location of these cells within the crypt in our cyclic IF images in an unbiased manner (**Supplementary fig. 4a**). Briefly, we trained HaloAI DenseNET V2 classifiers to distinguish crypt base (CD44+ and EpCAM+) and crypt top (CD44- and EpCAM+) epithelium and used Halo HighPlex FL v4.2.14 analysis to identify P14+ T cells based on CD3, CD8b, and Thy1.1 positivity. A Halo infiltration analysis was then performed to calculate the number of P14+ T cells infiltrated into the crypt top and crypt base epithelium, per millimetre squared of tissue (**Supplementary fig. 4b-c)**. Prior to antigen induction, endogenous T cells were evenly distributed between the crypt base and the crypt top epithelium (**Fig. 3e**). By contrast, P14 CD8 T cells were significantly more associated with the crypt base than the top in mice with NINJA induction. Thus, despite neoantigen expression throughout the crypt, neoantigen-specific CD8 T cells were preferentially found at the base of neoantigen+ crypts, where both Nur77 WT and Nur77 reporter P14 T cells can be observed (**Fig. 3e, Supplementary fig 4d**). Importantly, all Thy1.1+ CD45.1+ cells (Nur77 WT P14 T cells) were GFP-, as expected (**Supplementary fig 4d**).

### GP33-specific CD8+ T cells eliminate epithelial stem cells

The location of the infiltrating cytolytic CD8 T cells prompted investigation into whether these cells were killing the LGR5+ stem cells, which are known to reside at the base of the crypt^38, 39, 40^. To assess this, we sorted NINJA+ and NINJA-EpCAM+ epithelial cells from T cell-sufficient and T cell-depleted CNC mice at day 9 post-antigen induction, as we reasoned that this was a timepoint when the elimination of the epithelial cells would be occurring (**Supplementary fig 5a**). We then subjected these sorted epithelial cells to single-cell RNA sequencing using the 10x platform to assess which cell types were present in these four samples (**Fig. 4a**). We obtained transcriptomic data from 26,000 cells (4,501 NINJA+ T cell sufficient; 5,007 NINJA-T cell sufficient; 10,169 NINJA+ T cell deficient; 6,323 NINJA-T cell deficient). Analysis with Seurat identified 7 clusters that we projected onto 2-dimensional Uniform Manifold Approximation and Projection (UMAP) dimensionality reduction plots (**Fig. 4b**). We identified the clusters as LGR5+ Stem cells, transit amplifying cells, goblet cells, tuft cells, enteroendocrine cels, early enterocytes, and late enterocytes (**Supplementary fig. 5b, Supplementary Table 1**). Notably, the stem cell cluster was highly enriched in genes from a stem cell gene module, including high expression of Lgr5, Ascl2, CD44, Hopx, and Sox9 (**Supplementary Table 2**).

**Fig. 4:**
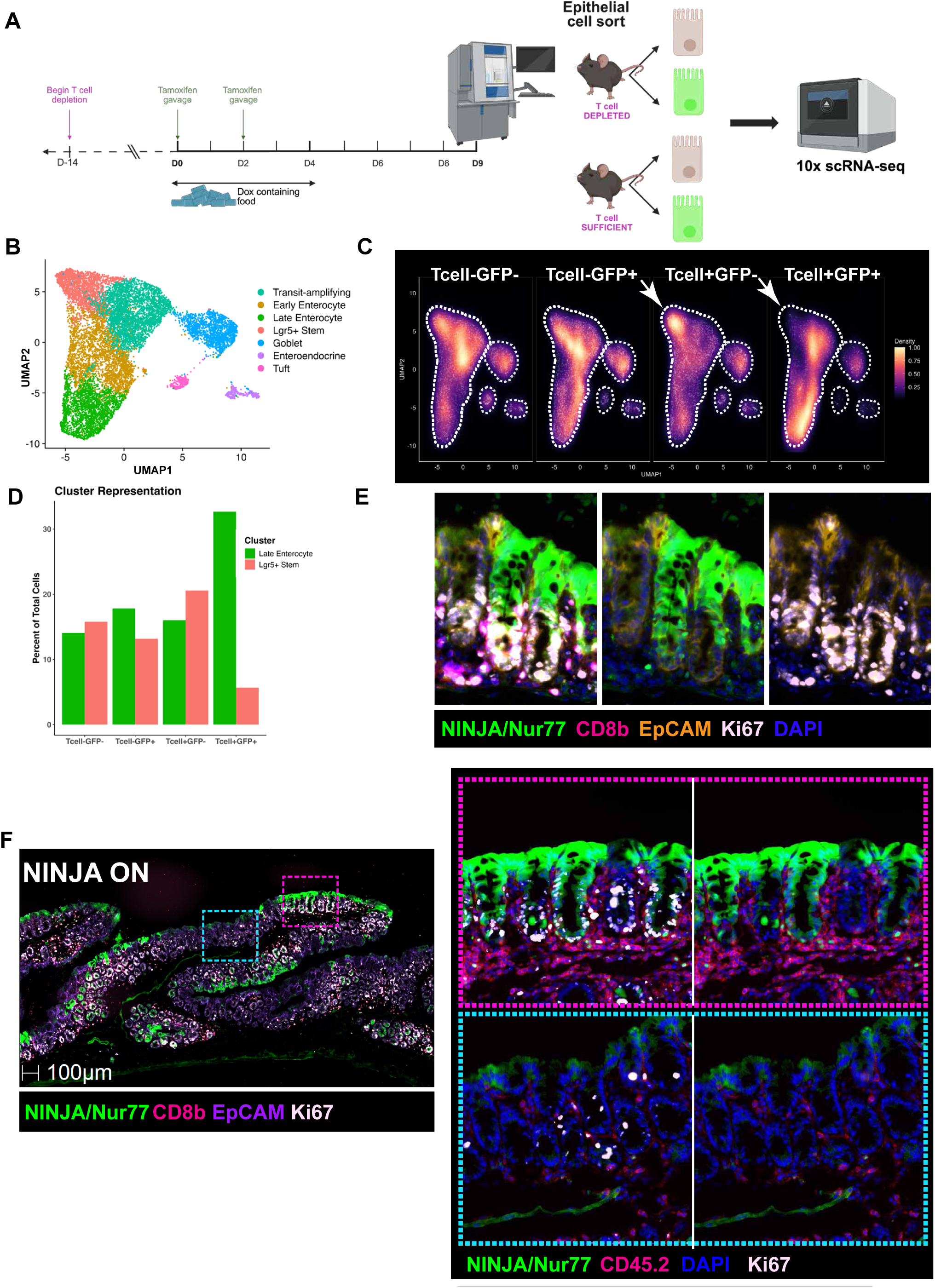
NINJA-specific T cells target LGR5+ stem cells. **a**, Experimental timeline of the scRNA-seq **b**, UMAP clustering and cell type definitions. **c**, Density plots of NINJA+ and NINJA-epithelial cells within the T cell-depleted and T cell-intact groups. The white arrows indicate the LGR5+ stem cluster location. **d**, Percent of cells in each condition in the LGR5+ stem cell and late enterocyte clusters. **e**, representative (n = 6) 20x CellDIVE immunofluorescence image from days 6-8 depicting Ki67 at the crypt base, green = NINJA and Nur77, white = Ki67, pink = CD8b, orange = EpCAM. **f**, representative 20x CellDIVE immunofluorescence image from days 6-8 (n = 6) depicting Ki67 near regions with NINJA+ expression and CD8b+ T cells, green = NINJA and Nur77, white = Ki67, pink = CD8b or CD45.2, blue = DAPI, purple = EpCAM. Blue dashed line is a NINJA negative region, and the pink dashed line is a NINJA expressing region.

Comparing the NINJA+ and NINJA-epithelial cells under T cell-deficient conditions, the distribution of cells across all 7 clusters was relatively similar (**Fig. 4c, Supplementary fig. 5c**). This suggested that the induction with CDX2-CreER had induced neoantigen across all major cell types in the colon, including stem cells, in a relatively unbiased manner. MHC class I was also expressed by the cells in all clusters (**Supplementary fig. 5d**), suggesting that all epithelial cell populations would be capable of presenting neoantigens to neoantigen-specific CD8 T cells. In the context of T cell sufficiency, however, there was a specific loss of NINJA+ epithelial cells in the stem cell cluster and an enrichment in late (differentiated) enterocytes, with NINJA-epithelial cells being more enriched in the stem cluster (**Fig. 4c, d**). Projection of the cellular distribution of the NINJA+ epithelial cells from T cell-sufficient and T cell-deficient conditions (density plot) also showed this selective depletion of cells with the stem-cell gene module (**Fig. 4c**). Together, these data indicated a selective loss of NINJA+ stem epithelial cells in the presence of T cells.

Despite the rapid loss of stem cells in the colon, mice did not exhibit signs of pathology (**Fig. 1f, Supplementary Fig. 1c**), suggesting a potential repair process. Others have observed that the loss of LGR5+ stem cells does not cause disease in the colon because Krt19+ epithelial cell populations rapidly replace them^41, 42, 43^. Immunofluorescence for Ki67 showed increased staining in crypts where NINJA+ cells were at the top but not the base(**Fig. 4e**). Additionally, there was increased epithelial Ki67 staining in regions of the colon with immune infiltration and NINJA expression (**Fig. 4f**), suggesting these cells were proliferating in response to CD8 T cell-driven elimination of the GFP+ stem cells. In line with this, we observed increased expression of Krt19 mRNA amongst the GFP-negative stem cells from mice with T cells (**Supplementary fig. 5e**), which we posit is the cell population that is enriched with repopulating stem cells. Together, these data suggest that NINJA+ stem cells in the colonic crypt epithelium have been eliminated, and the NINJA+ epithelial cells at the top of the crypt are being lost via extrusion into the gut lumen as part of the normal cycle of epithelial turnover.

### Mouse and human intestinal epithelial cells have an impaired ability to upregulate PD-L1

Our prior studies on T cell tolerance in the skin had demonstrated that GP33-specific CD8 T cells were restricted from causing pathology and eliminating NINJA+ epithelial cells through the PD-1/PD-L1 pathway ^29^. This process involved the post-transcriptional repression of effector proteins like Granzyme A, IFNg, and TNFa. In the colon, we also observed that GP33-specific CD8 T cells in the IEL expressed high amounts of PD-1. These PD-1+ CD8 T cells upregulated Granzymes A and B in the tissue and interacted directly with NINJA+ epithelial cells (**Figs. 2e-f and 3a**). We also noted that while NINJA+ crypts were surrounded by PD-L1+ myeloid cells, the NINJA+ epithelial cells were surprisingly PD-L1-negative (**Fig. 5a, Supplementary fig. 6a**; Note, EpCAM staining is shown to demonstrate staining for a membrane protein on epithelial cells). Previous work has established that pancreatic b cells upregulate PD-L1 in response to T cell infiltration ^44, 45^, so as a positive control, we activated NINJA on β cells in islets and observed upregulation of PD-L1 (**Supplemental fig. 6b**). We hypothesised that upregulation of Granzymes and perforin by GP33-specific CD8 T cells in the LP and IEL was due to antigen recognition in the absence of PD-L1. To test this, we transduced previously activated P14 CD8 T cells with either WT PD-1 or a signalling-dead mutant PD-1 (NULL PD-1) and then re-exposed them to antigen in the context of PD-L1-Ig (**Fig. 5b**). In the absence of PD-1 signals or PD-L1 protein (Fc control), we observed high amounts of cytolytic proteins. However, PD-L1 significantly suppressed the expression of perforin and Granzyme B at the protein level. Thus, in line with prior work, the cytolytic capacity of GP33-specific CD8 T cells was directly regulated by exposure to PD-L1. Moreover, the absence of PD-L1 was likely enabling elimination of neoantigen+ epithelial cells. Therefore, we next asked why these cells were not upregulating PD-L1.

**Fig. 5:**
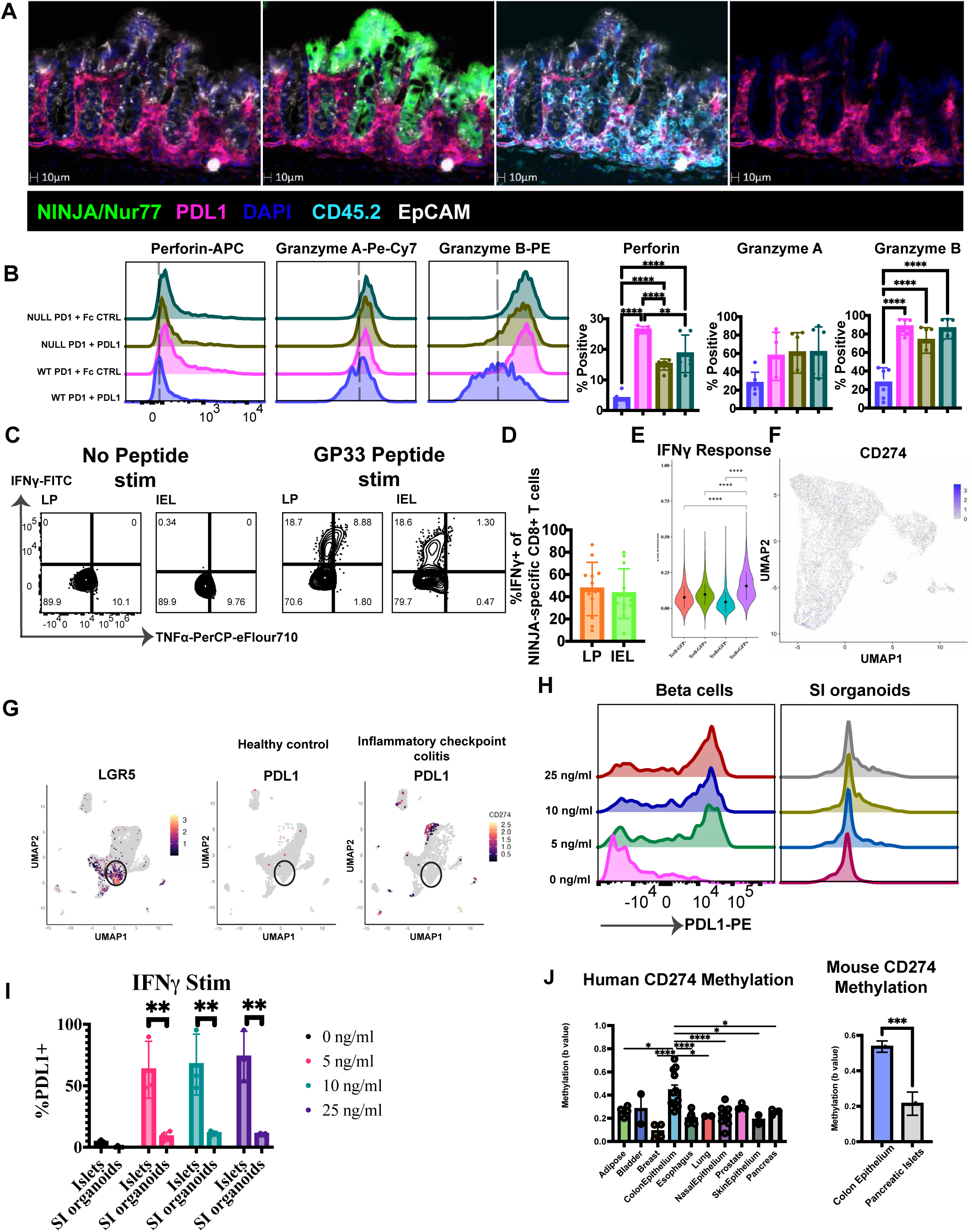
Epithelial cells have a blunted ability to upregulate PDL1. **a**, Representative (n = 6) 20x CellDIVE immunofluorescence image from days 6-8. GFP = NINJA and Nur77, pink = PDL1, blue = EpCAM, cyan = CD45, white = EpCAM. **b**, 48 hour in-vitro stimulation with CD3, CD28 and either PDL1 or an Fc control. Representative histogram expression of Perforin, Granzyme A, and Granzyme B with quantification of the percent expression. **c**, Ex-vivo GP33 peptide stimulation. Representative IFNγ and TNFα flow plots, pre-gated on CD8+ T cells. **d**, Quantification of the percent of antigen-specific T cells (GP33 and GP34 tetramer-positive CD8+ T cells) that make IFNγ following GP33 stimulation. **e**, KEGG IFNγ response module enrichment score. **f**, CD274 transcript expression. **g**, UMAP reduction with LGR5 expression and PDL1 expression from healthy control and inflammatory colitis conditions. **h**, representative histogram of PDL1 expression following a 24-hour in-vitro stimulation of primary human β cells and human small intestine organoids with various doses of IFNγ. **I**, Quantification of the %PDL1+ human β cells and human small intestine organoids following the IFN γ stimulation. **J**, Human CD274 promoter methylation across various organs and cell types and mouse CD274 promoter methylation in islets and colonic epithelial cells, three mice each, pooled with two technical replicates. To assess statistical significance of the PDL1 stimulations, a one-way ANOVA with Tukey’s multiple comparison test was performed. To assess the statistical significance of the KEGG IFNγ signature, Wilcoxon sign-rank tests were performed. To determine the statistical significance of the in-vitro IFNγ stimulations, a Mixed-effects model (REML) with Šídák’s multiple comparisons test was used. For all statistical tests,*P < 0.05; **P < 0.01; ***P < 0.001; ****P < 0.0001.

IFNg is a known inducer of PD-L1 expression on most cell types, and so our initial analyses focused on whether infiltrating GP33-specific CD8 T cells in the IEL and LP were able to make IFNg^45, 46^. For this, we utilised *ex vivo* peptide stimulation with GP33-43 peptide, followed by intracellular staining for the cytokines IFNg and TNFa (**Fig. 5c,d**). Ex vivo, CD8. T cells from both sites could make IFNg when stimulated with peptide, and IFNg+ CD8 T cells did not co-produce TNFa. This data suggested that GP33-specific CD8 T cells had the capacity to produce IFNg locally. However, to assess whether NINJA+ epithelial cells were seeing IFNg, we analysed the epithelial cell populations in our scRNAseq for the expression of the KEGG IFNg signalling gene (ISG) module (**Fig. 5e**). NINJA+ epithelial cells in T cell-sufficient animals had significant upregulation of the ISG module compared to NINJA-epithelial cells from the same animals. Moreover, this upregulation was T cell dependent. Yet, despite expression of a strong ISG signature, we were unable to detect expression of *Cd274* (gene for PD-L1) by any epithelial cell subset in our scRNAseq dataset (**Fig. 5f**). To validate that this was not a consequence of our model, we analysed primary C57Bl/6 epithelial cells for *Cd274* expression (**Supplementary Fig. 6c**). We were able to detect *Cd274* within differentiated cells, such as tuft cells, but not in epithelial stem cells.

Two questions arose from the above analyses: (1) whether similar results would be seen under conditions of chronic IFN signalling, and (2) in the context of overt intestinal inflammation in humans. To address the first, we analysed single-cell RNAseq data sets from a recent study that described an important role for IFNg signalling in the maintenance of aged mouse intestinal epithelial cells^47^. We observed an upregulation of interferon-stimulated genes (ISG) in the aged epithelium, but did not observe *Cd274* expression (**Supplementary fig. 7a-e**). To assess whether similar results were seen in the context of human colon during chronic inflammatory conditions, we reanalysed a dataset of single-cell RNAseq on colonic epithelial cells from patients with either active ulcerative colitis (UC) or checkpoint immunotherapy-induced colitis (CPI-colitis). This dataset featured controls without the inflammatory conditions and healthy controls, allowing us to assess changes in CD274 expression across a variety of physiological states^48^. Analyses of the patients with CPI-colitis revealed an increased ISG signature in epithelial stem cells compared to CPI-treated controls (**Supplementary fig. 8a-c**). Moreover, there was a fraction of CD274+ epithelial cells in CPI-colitis, called inflammatory enterocytes, that are more differentiated than stem and transit amplifying cell populations (**Fig. 5g**). These findings are in line with a recent study showing that macrophages and some crypt top epithelial cells were expressing PD-L1 in active CPI-induced colitis, but not epithelial cells at the crypt base^49^. We further found that CD274 was not detected in any of the epithelial cell populations from patients with active or inactive UC (**Supplemental Fig 8d**). Together, these data demonstrate that PD-L1 is not expressed by colonic epithelial cells in various settings of increased inflammation.

The above data suggested to us that perhaps the intestinal epithelium was resistant to upregulation of PD-L1 following IFNg stimulation. To test this, we challenged human small intestine epithelial organoid samples with varying concentrations of IFNg overnight and analysed PD-L1 upregulation the next day by FACS (**Fig. 5h, i**). As a positive control, we cultured human β cells overnight with IFNg. As expected, incubation with as little as 1ng/mL of IFNg led to robust upregulation of PD-L1 by beta cells, but even in the presence of 25 ng/mL, intestinal epithelial cells were mostly PD-L1-negative. Thus, intestinal epithelial cells were refractory to upregulation of PD-L1 in the context of IFNg exposure.

One of the primary mechanisms of regulation for *Cd274* in cancer cells is through hypermethylation of the *Cd274* promoter region. Analysis of human tissue methylation atlases showed that methylation of the promoter for *CD274* was significantly increased in colon epithelial tissue compared to several other tissues, including Adipose, Breast, Esophagus, Lung, Nasal, Skin epithelium, and Pancreas tissues (**Fig. 5j, Supplementary fig. 9a**). This increase was restricted to the promoter regions (**Supplementary fig. 9b**). To confirm that the Cd274 promoter in mouse was also methylated, we isolated epithelial cells from mouse proximal colon and cells from the pancreatic islets and performed genome-wide methylation sequencing (**Fig 5j**). This analysis also showed a significant increase in the methylation of the *Cd274* promoter in colon epithelial cells, consistent with the idea that the inability of colon epithelial cells to upregulate PD-L1 was due to a natural epigenetic silencing of the promoter for this gene.

## Discussion

The gut is a unique environment where tolerance is essential for preventing T cell-mediated pathology, yet effective immunosurveillance is necessary. This raises a clear question: how do both processes coexist to maintain homeostasis? Here, we established a model of immunosurveillance in which an antigen is inducibly expressed in the proximal colon epithelium, simulating a somatic mutation occurring in epithelial cells. We found that CD8+ T cells were primed in the colon-draining mesenteric lymph nodes and specifically infiltrated antigen-expressing crypt bases. Elimination of the proximal colon epithelial cells was associated with epithelial cell proliferation, replacing neoantigen+ epithelium with neoantigen-negative epithelium. PD-1+ CD8 T cells gained cytolytic capabilities in the tissue and directly killed epithelial stem cells via a perforin-dependent mechanism. Yet, despite a strong IFNg signature, antigen-expressing stem cells did not significantly upregulate PD-L1, which mirrored findings seen under other conditions of inflammation, like ageing and CPI-induced colitis. The epithelial cells in human and mouse exhibited hypermethylation of the PD-L1 promoter, suggesting an underlying mechanistic explanation. Additionally, stimulation in the presence of PD-L1 suppressed the expression of cytolytic molecules on PD-1+ CD8 T cells. Thus, in the absence of epithelial PD-L1, CD8 T cells were able to efficiently survey for and kill neoantigen+ epithelial cells in the proximal colon.

Previous studies of CD8 T cell responses to peripheral antigens have often relied on expression of a model antigen under a tissue-specific promoter. This literature suggested that tolerance was mediated via the priming phase; a T cell primed against an antigen without accompanying costimulation would be rendered tolerant in the LN by either deletion or anergy. For example, transferred OVA-specific CD8+ T cells responding to OVA expressed under the intestinal fatty acid binding protein iFABP failed to make much IFNγ upon *ex vivo* stimulation, and there was no overt immunopathology unless mice were also treated with vesicular stomatitis virus (VSV) encoding OVA ^50, 51^. Ova in iFABP-Ova mice was presented by DCs in the mLN, but also was expressed by fibroblastic reticular cells in the mLN stroma, and the latter was sufficient to drive CD8 T cell anergy/deletion in a PD-1 dependent manner ^52, 53, 54^. Depletion of FoxP3+CD8+ Tregs also led to pathology in the intestine of iFABP Ova mice ^55^. Yet, notably, in the tolerant state, Ova-specific CD8 T cells migrated into the intraepithelial layer of the small intestine. These infiltrating CD8 T cells were initially Granzyme B+ and responded to antigen in the tissue, but did not mediate immunopathology ^56, 57^. Indeed, the absence of overt pathology demonstrated that CD8 T cells in the intestine were tolerant, but there was no marker for elimination of antigen+ epithelial cells in this model. Remarkably, more recent studies demonstrated that challenge of iFABP-Ova mice with 3 rounds of Ova-expressing pathogens breaks tolerance, resulting in large numbers of functional Ova-specific effector CD8 T cells in the intestine ^58^. Yet, these mice still did not have intestinal pathology. In the CNC model, there was unambiguous CD8 T cell-driven epithelial cell elimination without pathology. This was only detectable because target cells were GFP+ and the crypts were replaced by GFP-negative epithelial cells. Notably, the fate of neoantigen+ cells in the colon differs from our findings in the skin, where GFP+ cells are maintained for 30+ days^29^. Thus, the biology of immunosurveillance appears distinct in the intestinal tissue.

Our findings on intestinal repair are reminiscent of studies that eliminated LGR5+ stem cells, either by using LGR5-DTR to kill stem cells in the colon/small intestine or using GFP-specific CD8 T cells (from JEDI mice) to kill GFP-expressing LGR5+ epithelial cells ^41, 43, 59^. In both cases, the replacement by proliferating epithelial cell populations masked the effects of stem cell elimination. In the small intestine, irradiation has been used to disrupt repair after elimination of LGR5+ stem cells. In the colon, this repair is mediated by a radio-resistant Krt19+ cell that resides above the LGR5+ stem-cell population ^41^. We posit that a similar cell is repopulating the crypt base after CD8 T cells eliminate neoantigen+ LGR5+ stem cells in CNC mice. In line with this, we see upregulation of *Krt19* amongst neoantigen-negative stem cells from mice where T cells are killing neoantigen+ cells (**Supplementary fig. 5e**). Notably, however, this repair of the tissue obscures the immunosurveillance function of the CD8 T cells in the tissue: the process is subclinical.

PD-1/PD-L1 interactions are central to the regulation of CD8 T cell responses across infection, autoimmune disease, cancer, and tolerance. In cancer and viral contexts, PD-1-PD-L1 interactions lead to decreased cytotoxicity, Perforin, and Granzyme B in effector CD8 T cells ^60, 61, 62, 63, 64, 65, 66, 67, 68^. PD-1 is expressed by most tissue-resident CD8 T cells^69^, so tissue upregulation of PD-L1 is key for suppression of immunopathology. In line with this, loss of PD-L1 by parenchymal cells accelerates the onset and severity of many autoimmune diseases ^70, 71, 72, 73^. Notably, PDL1 expression, specifically on parenchymal cells, abrogates diabetes induction caused by the transfer of activated OT-I cells into RIP-Ova mice, ^74^ which is in line with studies in the non-obese diabetic mice ^44, 70, 72^. Thus, PD-L1 expression on parenchymal cells potentially serves as the line of last defence for self-reactive CD8 T cells after infiltration of a critical site. The blunted ability of colonic epithelial cells to upregulate PD-L1 would ensure that they cannot protect themselves from killing by neoantigen-reactive CD8 T cells.

Analysis of tissues taken from patients with immune-checkpoint induced irAEs has shown upregulated expression of PD-L1 by tissue parenchymal cells in the heart, pancreatic islets, kidney, and other tissues ^44, 75, 76, 77^. By contrast, PD-L1 is only expressed by macrophages and crypt top epithelial cells in patients with CPI-colitis^49^. Furthermore, the colon is an uncommon site for irAEs following PD-1/PD-L1 blockade, the skin, another barrier tissue ^78^. PD-L1 has also been shown to play a role in protection from DSS-induced colitis^79^, but studies have observed that PD-L1 was expressed by myeloid cell populations and not intestinal epithelial cells ^54, 80^. Data from cancer suggest that PD-L1 induction can be regulated epigenetically, with alterations in DNA accessibility and DNA methylation in the promoter as key determinants. Hypomethylation has been consistently correlated with higher PD-L1 mRNA expression across tumour types, and short-term treatment with the hypomethylating agent 5-azacytidine led to increased PD-L1 expression ^81, 82, 83, 84, 85^. We found that colon epithelium has high methylation of the PD-L1 promoter in both human and mouse epithelium, but this was not true of other regions of the gene, suggesting specificity. The methylation status of this locus likely changes during the progression of colon cancer, which would serve to aid tumours in their development.

When contemplating immunosurveillance nearly 75 years ago, Macfarlane Burnett envisioned immune cells surveying for “foreign” antigens, with somatic mutation yielding detectable antigens that could distinguish pre-cancerous cells ^86^. Yet, he also recognised the enormous challenge of finding these rare mutant cells within healthy tissues. This challenge is exemplified by the nature of the colon: there are ∼10-15 million crypts in the adult human colon, each of which is comprised of 2000 cells ^87, 88^; thus, effective immunosurveillance *would* require CD8 T cells to screen 20-30 billion epithelial cells. Burnet noted that mutations in “germ cells” would lead to more progeny, but also that “a mutation in a cell giving rise to highly expendable descendants may have wider influence” ^86^. Immunosurveillance, he concluded, would require expansion of clones of mutant cells in the tissue, which would be provided as a consequence of tumorigenesis. In the absence of tumorigenesis, the dynamics of stem cells in the colon provide a means for expanding mutations. Neutral and selected drift results in an ancestral clone age of ∼5.5 years among the ∼7 stem cells/crypts. Consequently, in the healthy human colon, many of the 1,500-15,000 somatic mutations identified are shared among all the cells of each crypt^11, 14, 16^. This process fixes the mutations into the crypt, while rare crypt fissions expand the influence of mutation to neighbouring crypts ^89^. Thus, over decades of life, somatic mosaicism results throughout the colon tissue. We posit that the ability of CD8 T cells to focus on stem cells has evolved as a means for assessing the neoantigen burden of the crypt. Coupled with natural processes that increase representation of stem cells with potentially harmful mutations, this would provide an effective surveillance mechanism for clearing the most dangerous stem cells, without causing broader immunopathology that would be detrimental to the host.

## Limitations

There are limitations to our study. NINJA relies on doxycycline and tamoxifen for antigen induction, which may impact the immune response. Additionally, the doxycycline is given in the chow. Thus, the dosing and dosing schedule will vary between mice, meaning variable kinetics and antigen burden. Furthermore, the requirement of a Cre means that antigen is induced in many crypts at once, making it challenging to test different doses of antigen. There are also still open questions. We do not know how the T cells are able to retain function after priming without inflammation as a source of costimulation. Additionally, we do not know how the T cells are able to specifically target the crypt base. However, recent work has suggested that it may be due to chemokine gradients within the crypt and the location where the T cells enter the tissue ^90, 91^.

## Methods

### Mice

All studies were carried out in accordance with the guidelines of the Declaration of Helsinki and approved by the Institutional Animal Care and Use Committees of Yale University (IACUC no. 2025_20112). All mice were bred in specific pathogen-free conditions. Six-to sixteen-week-old female and male mice were randomly allocated to experimental groups. Mice were housed with enrichment nestlets in a controlled environment with ad libitum Access to food and water. There was a fixed 12-hour light cycle at an ambient temperature of ∼23°C. NINJA mice (066733, JAX) were bred to CAG-rtTA3 mice (016532, JAX) and referred to as NINJA. Our P14 mice are bred in-house and were crossed to Nur77-eGFP mice (018974, JAX)

### NINJA Induction

Mice were provided with ad. Lib. access to doxycycline hyclate (625 mg/kg) containing food (Inotiv, TD.120769) from days 0 to 4. Tamoxifen (220 μl, 20 mg/ml in corn oil, MP biochemical) was gavaged orally on days 0 and 2.

### TdTomato Induction

CDX2CreERT2 x Ai14 mice were gavaged orally with tamoxifen (220 μl, 20 mg/ml in corn oil, MP biochemical) on days 0 and 2.

### Tissue processing

Lymph nodes were smashed through 70-micron filters using FACS buffer (2% FBS, 1mM EDTA in PBS w/out CA^2+^ and Mg^2+^).

For epithelial cells, a modified protocol based on Gracz AD, Puthoff BJ, and Magness ST (2012) was used. Briefly, harvested colons were cut longitudinally and cleaned by shaking in ice-cold DPBS. Then, colons were incubated in a 50 ml conical containing 10 mL of ice-cold DPBS, 10 mM EDTA, 30 mM DTT, 10 μM Y27632 (Tocris, 1254) for 20 min on ice, followed by a 10 min incubation at 37**°**C in 6 mL of DPBS, 10 mM EDTA, 10 μM Y27632. Next, the tube was manually shaken for 45s to remove the epithelial cells. After, the tissue was discarded from the tube, and the remaining solution was pelleted by centrifugation at 1000g for 5 min at 4**°**C. The pellet was resuspended in 10 mL of prewarmed magnesium and calcium-containing HBSS with 0.8 mg/mL dispase II (ThermoFisher Scientific, 17105041). The conicals were rotated at 37**°**C for 10 min, followed by a quench with 10ml of 10% FBS. The cells were then sequentially filtered through 100-, 70-, and 40-micron filters on ice. The cells were then pelleted at 4**°**C, 1000g for 5 min, and resuspended for downstream analysis such as flow cytometry or sorting.

For lymphocytes within the colon, a modified protocol graciously provided by the Hand lab was used. Briefly, harvested colons were cut longitudinally and cleaned by shaking in ice-cold DPBS. Then, the colons were placed in a 75 mL flask containing 20 mL of RPMI, 25 mM HEPES (ThermoFisher Scientific, 15630080), 3% FBS, 5 mM EDTA (Millipore Sigma, E7889-100ML), and 0.145 mg/mL of DTT (Millipore Sigma, 10197777001). The flasks contained a stir bar and were covered with foil. They were then placed on a stir plate for 20 min, 400 rpm, in a 37°C/5% CO_2_ incubator. The colons were placed in 25 mL conicals containing 15 mL RPMI, 25 mM HEPES, and 2 mM EDTA. To remove the epithelial cells, the conicals were vigorously shaken for 1 min. For lamina propria processing, the tissue was then placed in a 50 ml flask with 6 mL of prewarmed digestion media, consisting of RPMI containing 25 mM HEPES, Liberase TL (1:250 of 25 mg/ml stock, MilliporeSigma, 05401020001) + 0.05% DNAse I (MilliporeSigma, 11284932001). The flask was stirred, covered in foil, at 400 rpm in a 37°C/5% CO_2_ incubator for 15 min. Then, another 6 mL of digestion media was added for a 15-minute incubation. Each flask was quenched with 25 mL of 3% FBS RPMI and sequentially passed through 100-, 70-, and 40-micron filters on ice. The solution was pelleted at 400 g, 4°C, for 5 min, and resuspended for downstream analysis. To harvest from the epithelial layer, the epithelial cell-containing solution (without the tissue) from the vigorous shaking step above was sequentially moved through 70- and 40-micron filters on ice. Then, the solution was pelleted by 400 g, 4°C for 5 min. The cells were resuspended in 30% room temp Percoll (Cytivia, 17089101) and centrifuged for 20 min, 22°C, with a break set to 1. The Percoll was discarded, and the remaining cells in the pellet were resuspended for downstream analysis.

### Immunofluorescence

Samples were fixed overnight in paraformaldehyde-lysine-periodate (PLP) at 4°C. The samples were then placed in 30% sucrose for 6-24 hrs. Tissues were embedded in TissueTek O.C.T. compound and then placed on dry ice for freezing. Tissues were then sectioned to 4-7 µm using a cryostat. The tissue sections were processed for imaging on the Cell DIVE (Leica Microsystems), a platform for multiplex imaging.

Briefly, slides were rehydrated with 3 x 5 min PBS incubations. The sections were then incubated with 0.2 % BSA, 0.05% sodium azide, 0.3% Triton X-100, and 10% FBS for 1 hr for blocking and permeabilization. Slides were then placed into ClickWells (Leica Microsystems) and washed with 0.01% Tween-20 in PBS (2 x 5 min) PBST. Sections were then stained with 1 μg/ml DAPI for 15 min, followed by another wash with the PBST. 1 ml of 50% glycerol was added, and then the slides were imaged for the first autofluorescence round. Then, unless otherwise noted, the slides were stained with antibodies at 1:100 in 3% BSA for 1 hr. The slides were then washed with PBST (2 x 2 min) followed by DAPI staining for 2 min, and then another 2 min wash. 1 ml of glycerol was added for biomarker imaging. After the imaging, slides were washed with 3 x 5 min with PBST. The sections were bleached with dye inactivation solution (100 mM NaHCO_3_, 3% H_2_O_2,_ in ddH_2_O) for 15 minutes and then washed again with PBST (3 x 5 min). Slides were then stained for biomarkers, imaged, and then dye-inactivated for up to 8 rounds until the image was completed. Imaging workflow followed Leica Cell DIVE multiplex imaging solution, and image results were analyzed by HALO AI image analysis software (Indica Labs) The antibodies used were CD8b clone YTS156.7.7, PDL1 clone 10F.9G2 (1:200), CD3 clone 17A2 (1: 200), Thy1.1 clone OX-7, CD45.1 clone A20, CD45.2 cone 104, EpCAM clone G8.8, CD44 clone IM7, CD4 clone RM4-5, CD31 clone MEC13.3, PD-1 clone RMP1-30, CD11c clone N418, Ki67 clone D3B5, F4/80 clone BM8, anti-GFP (polyclonal, Thermo Fisher Scientific cat# A-11122).

### Image analysis

HaloAI (Indica Labs) was used to analyze CellDive images. DenseNetV2 neural network classifiers were trained to recognize epithelium, crypt bases vs tops, and NINJA expression. The classifiers were trained based on the expression of GFP, EpCAM, CD44, and DAPI until the cross-entropy score reached <0.1. A classifier pipeline of the three classifiers was used to annotate the slides. First, the epithelium was annotated. Then, of the epithelium, the NINJA-expressing epithelium was annotated. In the last step of the pipeline, NINJA+ crypt bases and crypt tops were annotated. Following the classifier pipeline, a HighPlex FL v4.2.14 analysis was run to identify P14s based on Thy1.1 and CD8b positivity. Finally, an infiltration analysis was completed, which quantified the number of P14s within NINJA + crypt bases and crypt tops, and then normalized the P14 number to the tissue area.

### In vivo antibody depletions

200 µg of depletion antibody was injected intraperitonially (200 µl, 1mg/ml) every three days throughout the entirety of the experiment, with five doses given before antigen induction. *InVivo*MAb anti-CD4 clone GK1.5, *InVivo*MAb anti-CD8a clone 2.43, and *InVivo*Mab anti-Thy1.2 clone 30H12 were purchased from BioXcell.

### Naïve CRISPR-Cas9 electroporation

Naïve P14 CD8 T cells were isolated from spleens using the EasySep CD8 T cell negative selection kit (STEMCELL Technologies, 19853) according to the manufacturer’s instructions. The gene-editing protocol was adapted from Nussing et al, 2020. Briefly, sgRNA (Synthego) targeting either Perforin (5’-GCCCAGGAGGAACAGGCACG-3’ and 5’-CGUCACGUCGAAGUACUUGG-3’ in combination) or NTC (5′-GCGAGGTATTCGGCTCCGCG-3′ and 5’-CCAACCCGGCATCGTCCGCT-3’ in combination) was incubated with Cas9 nuclease V3 (IDT, 1081059) to create CRISPR-Cas9 ribonucleoprotein (RNP). In total, 1 × 10^5^ P14 CD8 T cells were electroporated with Perforin or NTC Cas9–sgRNA RNPs using the Lonza P3 Primary Cell 4D-Nucleofector X kit (V4XP-3032), according to the manufacturer’s instructions, and injected retro-orbitally into mice.

### Flow cytometry and in vitro antigen-specific stimulation

Flow cytometry analysis was performed on a BD Symphony, BD LSRII, Beckman Cytoflex LX, and Cytek Northern Lights. Cells were harvested from various tissues as described above. For in vitro stimulations, cells were cultured for 6 h at 37 °C in 5% CO_2_ in complete RPMI-1640 (10% HI-FBS, 55 μM beta-mercaptoethanol, 1x penicillin-streptomycin (ThermoFisher Scientific, 15-140-163), and 2 mM glutamine (ThermoFisher Scientific, 25030081)) supplemented with Brefeldin A (ThermoFisher Scientific, 00-4506-51) and either LCMV GP_33–41_ peptide (0.5 µg ml^−1^, AnaSpec, AS-61296) or left unstimulated in the presence of 5 × 10^5^ RAG KO splenocytes.

Cells were stained extracellularly with antibodies listed in **Supplementary table 3**. Staining was completed 1:200 in FACS (2% FBS, 1mM EDTA, in PBS w/out CA^2+^ and Mg^2+^) buffer at 4°C, protected from light. For intracellular staining, cells were processed using the Cytofix/Cytoperm kit (BD, 554714) following the manufacturer’s instructions, with a modification of staining intracellular antibodies overnight.

### IFNγ stimulations

The human islet processing and stimulation protocol was adapted from Perdigoto et al. (2022. Human islets from healthy, non-diabetic donors were obtained from Prodo Labs (Aliso Viejo, CA). The Islets were dissociated with a 10-minute TrypLE Express incubation in a 37°C bead bath and subsequently quenched with complete CMRL 1066 media (10% FBS, 10 mM HEPES, and 2 mM L-glutamine) (ThermoFisher Scientific, 11530037). Disassociated islets were incubated overnight at 37°C in 5% CO_2_ in complete media supplemented with the indicated amounts of IFNγ (R&D Systems, 285-IF-100/CF).

For human small intestine organoids, crypts were harvested from cryopreserved small intestine samples from resected healthy tissue. Organoids were cultured using STEMCELL Technologies’ Human IntestiCult Organoid system according to the manufacturer’s instructions. Organoids were treated with indicated amounts of IFNγ supplemented in the IntestiCult Organoid media overnight.

The IFNγ used for all treatments, for both the Islets and the small intestine organoids, came from the same lot.

### PDL1 in vitro stimulation

P14 T cells were isolated from splenocytes using the EasySep CD8 T cell negative selection kit (STEMCELL Technologies, 19853) according to the manufacturer’s instructions. T cells were then activated with anti-CD3/CD28 Dynabeads (ThermoFisher Scientific, 11452D) at a 1:1 ratio in the presence of recombinant human IL-2 at 5 ng/mL (Peprotech 200-02). On day 1 of activation, T cells were spinfected with retroviral supernatant in the presence of protamine sulfate (5 μg/mL) at 600g for 90 min. The retrovirus contained either a PD-1 overexpression construct or a PD-1 construct unable to signal (PD-1 NULL) because of mutated ITIM and ITSM (Y to F) signalling domains. On day 6 after spinfection, 1 x 10^5^ P14s were plated in 96 well plates coated with 3 µg/ml CD3 and 3 µg/ml CD28, with either recombinant 20 µg/ml PDL1 Fc chimera protein (R&D Systems, 1019-B7-100) or 10 µg/ml (equal molar ratio) recombinant human IgG1 Fc protein as a control (R&D Systems, 10-HG-100). After a 24-hour incubation in 37°C/5% CO_2_, P14 T cells were harvested for analysis.

### DSS colitis

4.2% dextran sodium sulfate (colitis grade, MW 36,000-50,000 Da, MP Biomedicals, 02160110-CF) was added to autoclaved drinking water, and mice had free access. Humane endpoint was reached with 15% bodyweight loss.

### Histopathology analysis

Whole colons were immersion fixed in 10% neutral buffered formalin and submitted to the Comparative Pathology Research Core in the Department of Comparative Medicine, Yale University School of Medicine, New Haven, CT, for processing, embedding, sectioning, and staining for hematoxylin and eosin (HE) and periodic acid-Schiff (PAS) by routine methods. The slides were examined blind to experimental manipulation and scored by semiquantitative analysis modified for colon from heart as previously described ^92^. Slides were analyzed using an Olympus BX53 microscope, imaged and photographed using an Olympus DP28 Camera, and cellSans standard 4.2.1 software. Images were optimized using Adobe Photoshop 26.8.1

### LCMV infections

B6 mice were infected with LCMV Armstrong at 2 × 10^5^ plaque-forming units (pfu) per mouse intraperitoneally. Mice were euthanized on day 8 after infection to collect spleens. LCMV Armstrong was produced in-house.

### Photographs of TdTomato+ intestines

The Xite Portable Fluorescence Flashlight System (green, NIGHTSEA) was used to photograph intestines from CDX2CreERT2 x Ai14 mice, 6 weeks after TdTomato induction.

### PDL1 scRNA expression on C57BL/6 mice

#### Mouse intestinal and colonic crypts isolation

Intestinal or colonic crypt isolation was performed on C57BL/6J (strain name: 000664) mice from the Jackson laboratory as previously reported ^93, 94^. Briefly, the entire small intestine or colon was dissected, and external fat, connective tissue, and blood vessels were carefully removed. Tissues were flushed thoroughly with ice-cold 1× PBS to remove luminal contents, then opened longitudinally. The intestine or colon was cut into 3–5 cm segments and transferred into ice-cold 1× PBS containing EDTA. For the small intestine, 7.5 mM EDTA was used with gentle rocking at 4°C for 30 minutes. For colon, 10 mM EDTA was used, and incubation was extended to 1 hour under the same conditions. Following incubation, the tissue fragments were vigorously shaken to release epithelial crypts into suspension. The supernatant was passed through a 70 µm cell strainer to remove villi (for intestine) or residual tissue fragments (for colon). Isolated crypts were then washed with ice-cold PBS and collected by centrifugation at 300 × g for 5 minutes.

Intestinal epithelial cells (IECs) were isolated by dissociation of the crypt suspensions into single cells with TrypLE Express (Cat# 12604-013, Invitrogen). Dissociated single cells were labeled with an antibody cocktail containing EPCAM-APC (1:400, Cat# 17-5791-82, eBioscience, G8.8), CD24-PE-Cy7 (1:400, Cat# 25-0242-82, eBioscience), and CD45-Alexa fluor 488 (1:400, Cat# 12-0451-83, eBioscience). Dead cells were excluded from the analysis with the viability dye SYTOX (Cat# S34857, Life Technologies). IECs were isolated as Epcam^+^ CD45**^-^** SYTOX**^-^** with a BD FACS Aria II SORP cell sorter into a supplemented crypt culture medium for culture or TRIzol reagent (Cat# 15596018, Thermo Fisher) to perform gene expression analysis.

#### Single Cell RNA Sequencing Library Preparation

For single cell sequencing of the mouse intestinal crypts, single cells were pelleted, washed and resuspended in FACS buffer (1X PBS, 10 μM Y-27632, 1% FBS, 0.5 mM EDTA) and passed through a 100μm FlowMi cell strainer (Sigma). DAPI was used for viability assessment. DAPI-negative cells were sorted by Sony SH800S sorter and single cell droplets were immediately prepared on the 10X Chromium according to manufacturer instructions at Cold Spring Harbor Laboratory Single Cell Facility. Single cell libraries were prepared using a 10X Genomics Chromium Controller (Cat #120223, 10X Genomics) and the 10X Genomics Chromium Next GEM Single Cell 3’ Gene Expression kit (Cat #1000268, 10X Genomics) according to the manufacturer’s instructions. Cell suspensions were adjusted to target a yield of 8,000 cells per sample. cDNA and libraries were checked for quality on Agilent Bioanalyzer, quantified by KAPA qPCR, and sequenced on a NextSeq500 (Illumina) to an average depth of approximately 33,000 reads per cell. The Cellranger pipeline (v4.0.0 10X Genomics) was used to align FASTQs to the mouse reference genome (gex-mm10-2020-A, 10X Genomics) and produce digital gene-cell counts matrices with default parameters.

#### Single Cell RNA-Seq data analysis

Single cell datasets were assessed for data quality following guidelines described previously ^95, 96^. Quality control and filtering was performed as described by ^95, 96^. After quality control (QC), normalization, graph-based clustering, and differential expression analysis was performed in Seurat (v5.1.0^97^). Datasets were normalized with SCTransform and the 5000 most variably expressed genes were identified with SelectIntegrationFeatures^93^. For each individual experiment, conditions were integrated into a singular dataset using the PrepSCTIntegration, FindIntegrationAnchors, and IntegrateData functions^94^. Missing values were imputed and technical noise accounted for via MAGIC imputation^98^. Normalization, scaling and clustering was similarly performed in Seurat. Subsequently, UMAP analysis and clustering was performed with the runUMAP function using the top 30 principal components (PCs) identified by RunPCA. Clustering was performed using the Louvain algorithm at a resolution of 1 on a nearest neighbor graph constructed with the FindNeighbors function. Clusters were labeled in accordance with expression levels of intestinal cell subtype signatures identified by^99^. Wilcoxon rank-sum tests using the wilcox.test() function in stats (v4.1.0^100^) were performed to determine if gene expression changes were significant.

### CNC Single-cell RNA sequencing

NINJA+ and NINJA-epithelial cells were sorted using a Bigfoot Spectral Cell Sorter (Thermo Fisher Scientific) from the proximal colons of CNC mice, 9 days post-antigen induction, that were either T-cell-depleted or T-cell-intact. Each sample consisted of three pooled mice from each condition. For the 10x 3’ v3.1 scRNA library prep, each condition was loaded onto its own Chromium instrument lane. Library preps were sequenced on an Illumina Novaseq at a depth of 20,000 reads per cell. Library preparation and sequencing were performed according to the manufacturer’s instructions by the Yale Center for Genome Analysis (3′ v3.1 single-cell RNA sequencing, 10X Genomics) (YCGA), the same day as sample collection. Data were processed with Cell Ranger 7.1.0 using the mm10 reference genome.

### PDL1 Promoter DNA methylation

For human tissues, methylation data for normal tissues were obtained from the Production Encode epigenetic dataset (GEO accession PRJNA63443), integrated, and analysed using the ENCODE portal and uniform processing pipeline^101, 102, 103^. For DNA methylation analysis from murine tissues, cells were enriched. DNA was isolated using the Quick-DNA Microprep Kit (ZymoResearch), bisulfite-converted according to the manufacturer’s instructions, quantified using the Qubit Fluorometric Analyzer (Thermo Fisher Scientific), and 250 ng of DNA was hybridised to the Mouse Methylation BeadChip array by the Yale Center for Genome Analysis. Idat files were processed, normalised, and methylation values were calculated using the SesaME and minfi packages^104, 105^. Methylation levels were compared using ANOVA and Welch’s t-test.

### Quantifications and statistical analysis

FlowJo v10.10.0 software was used for flow cytometry analysis. Prism v10.2.2 software was used to determine statistical significance. *P* values and associated statistical tests are indicated in each figure legend. Data are presented as mean +/- SD unless otherwise noted.

## Supporting information

Supplemental table 1. scRNAseq cluster definitions

Supplemental table 2. scRNAseq cluster DEGs

Supplemental table 3. Antibodies

## Acknowledgements

We thank all Joshi lab members for their helpful discussions and for reviewing this manuscript. This work was supported by T32 AI155387 (JB) and in part by the American Cancer Society (135667/RSG-21-105-01-IBCD) and the Colton Center for Autoimmunity at Yale. S.B. was supported by NIH R37CA292807 and Mark Foundation for Cancer Research (20-028-EDV). N.S.J. was supported by the Mark Foundation Emerging Leader Award and the Chan Zuckerberg Biohub Investigator Award. We thank the Yale Center for Cellular & Molecular Imaging for the usage of their Stellaris 8 DIVE confocal microscope, which is in part funded by the Yale Liver Center Morphology Core (NIH grant P30DK034989). We also thank the laboratory of Dr. Timothy Hand (Univ. of Pittsburgh) for the colon preparation protocol, the Yale Flow Cytometry Core, and Yale School of Medicine Comparative Pathology Research Core. The Yale Flow Cytometry Core is supported in part by NIH grants P30CA016359 and S10OD026996.

## Declaration of completing interests

The authors declare no competing interests.

## Additional information

Source data are provided with this paper.

Supplementary Information is available for this paper.

Correspondence and requests for materials should be addressed to Nikhil S. Joshi.

---

**Supplementary fig. 1.**
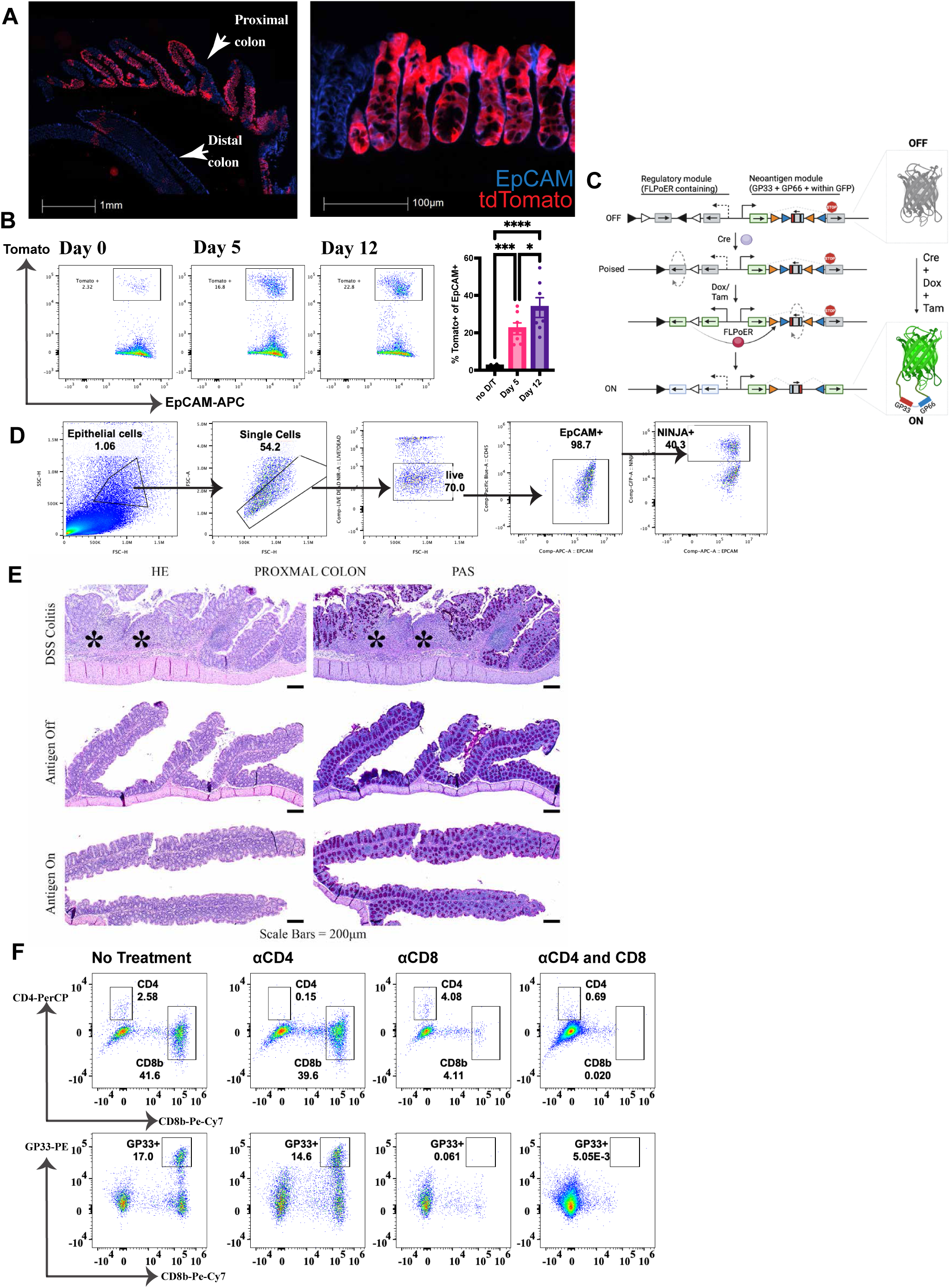
**a**, 20x CellDIVE immunofluorescence image depicting tdTomato within Cai14 mice 7 days post tdTomato induction. White arrows point to the proximal colon and the distal colon. blue = EpCAM and red = tdTomato. n = 4 mice, 1 experiment. **b**, Cai14 time course. Flow cytometry and quantification of tdTomato+ epithelial cells from no treatment (n = 8), day 5 (n = 7), and day 12 (n = 7) experimental groups. n is reported as the total across two experimental replicates. **c**, Schematic of NINJA allele and the recombination events in the regulatory module and neoantigen module required for NINJA expression. **d**, Gating strategy for epithelial cells. **e**, Representative images from H&E (Hematoxylin and Eosin)- and PAS (Periodic Acid Shiff)-stained sections of proximal mouse colon from DSS colitis control mice (n = 10) with loss of crypts, inflammation, and edema compared to unremarkable proximal colon from mice with antigen on (n = 9), antigen off (n = 10), and C57BL/6 treated with dox/tam (n = 12). n is reported as the total across three experimental replicates, except for the DSS group, which is two experimental replicates. **f**, representative flow cytometry plots depicting T cell depletion efficacy. Plots in the second row were pregated on CD8b, depicted in the row above. One-way ANOVA with Holm-Šídák’s multiple comparisons test was performed to determine the statistical significance of the tdTomato flow cytometry. *P < 0.05; **P < 0.01; ***P < 0.001; ****P < 0.0001.

**Supplementary fig. 2.**
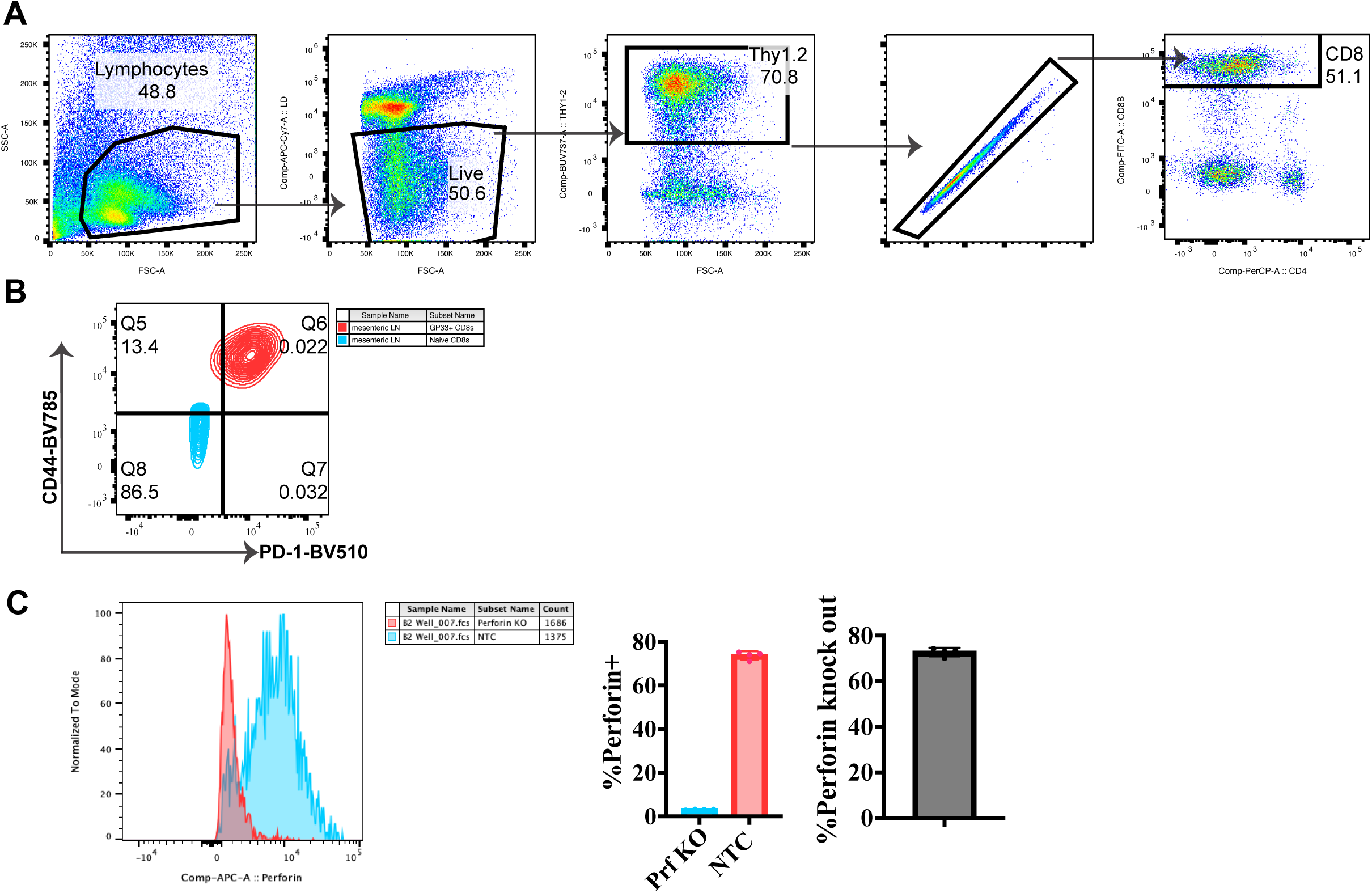
**a**, Flow cytometry gating strategy for CD8b+ T cells. **b**, Representative contour plot depicting CD44 and PD-1 staining on naive CD8b+ T cells (CD62L+ and CD44-) and GP33 specific CD8b+ T cells from the mesenteric lymph node. **c**, Validation of perforin CRISPR knock-out. Congenically marked naive WT and Perforin KO P14+ T cells were mixed 50:50 and adoptively transferred into C57Bl/6 mice, which were subsequently infected with LCMV Armstrong. On day 8, P14+ T cells were harvested from spleens. Perforin histogram staining for WT and PRF KO P14+ T cells with quantification of percent of P14+ T cells expressing perforin and knockout efficiency. Knockout efficiency was calculated as: (WT perforin expression - KO perforin expression)/WT perforin expression. n = 4, 1 experiment.

**Supplementary fig. 3.**
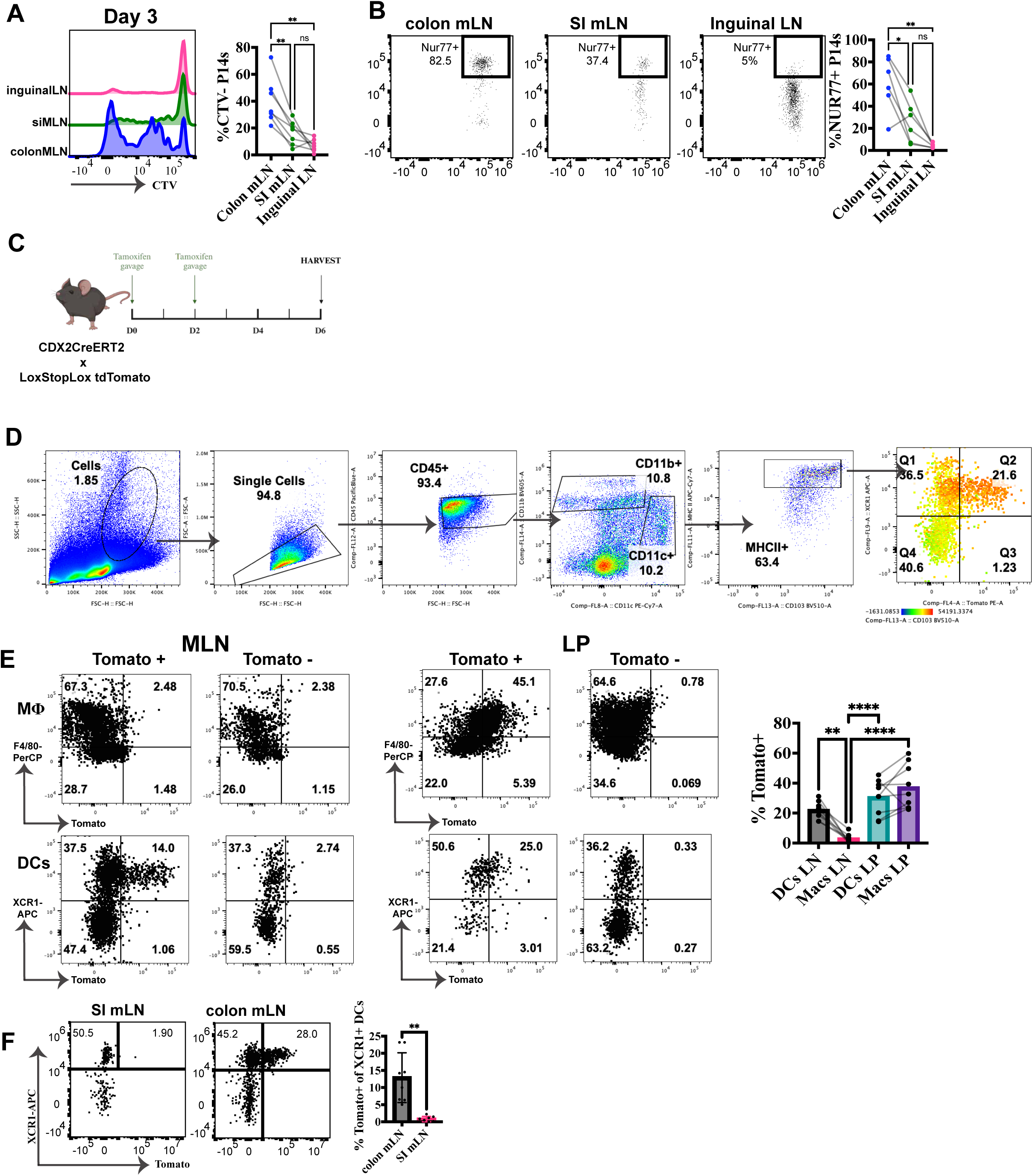
**a-b**, Cell trace violet (CTV) labelled Nur77 reporter P14+ T cells were adoptively transferred one day prior to antigen induction in CNC mice, consisting of two experimental repeats. **a**, Histogram depicting Day 2 CTV dilution (n = 6) in P14 T cells with quantification of the percent of P14 T cells with CTV dilution. Includes P14 T cells from the inguinal (control), proximal colon draining mesenteric, and distal ileum draining mesenteric lymph nodes. **b**, Representative dot plots and quantification of the frequency of Nur77+ P14 T cells (n = 7) from the inguinal (control), colon draining mesenteric, and small intestine draining mesenteric lymph nodes. **c**, Tracking of tdTomato uptake experiment schematic. **d**, Gating strategy for myeloid cells. **e**, Frequency of tdTomato within macrophages and dendritic cells in the mesenteric lymph node and lamina propria with representative dot plots. n = 9 for the lamina propria and n = 8 for the LN across three experimental replicates. **f**, Frequency of tdTomato within dendritic cells in the proximal colon draining mesenteric lymph node and the ileum draining mesenteric lymph node, with representative dot plots. n = 8 across three experimental replicates. A one-way ANOVA with Dunnett’s multiple comparison test was performed to determine statistical significance of the tomato frequency in myeloid populations within the LP and LN. A paired t-test was performed to determine the statistical significance of tomato uptake in dendritic cells between the ileum and proximal colon draining lymph nodes. *P < 0.05; **P < 0.01; ***P < 0.001; ****P < 0.0001.

**Supplementary fig. 4.**
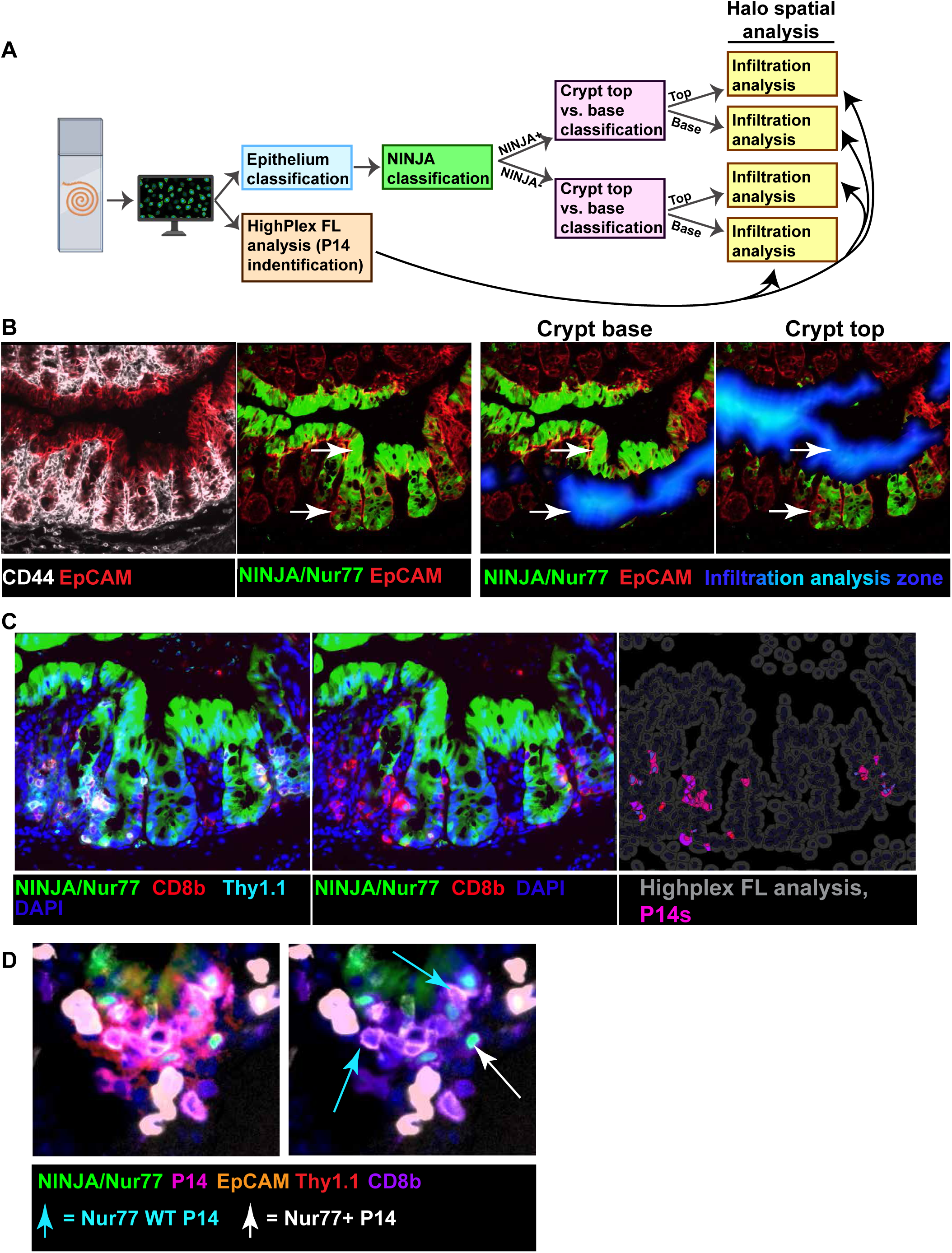
**a**,Schematic of Halo analysis pipeline. First, the epithelium classifier identifies the epithelial layer. Then, of the epithelial layer, NINJA+ and NINJA-epithelium is identified. Then, a classifier is used on both the NINJA+ and NINJA-epithelium to determine if it is in the crypt base or top. Additionally, a HighPlex FL analysis is performed on the entire tissue to identify P14+ T cells. Finally, an infiltration analysis determines the number of P14+ T cells within the NINJA+ and NINJA-, crypt base and crypt top epithelium, and normalises the number to the tissue surface area. **b**, 20x CellDIVE immunofluorescence images (green = NINJA and Nur77, red = EpCAM, white = CD44) and visualisation of the infiltration analysis area (blue) for NINJA+ crypt top and NINJA+ crypt base epithelium. **c**, 20x CellDIVE immunofluorescence images (green = NINJA and Nur77, red = CD8b, cyan = Thy1.1, and blue = DAPI) with HighPLEX FL P14+ T cell identification visual (pink and purple). **d**, 20x CellDIVE immunofluorescence image. Green = NINJA and Nur77, red = Thy1.1, purple = CD8b, pink (red + purple) = P14+ T cells (Thy1.1 + CD8b). Blue arrows indicate Nur77 WT P14+ T cells, and white arrows indicate Nur77 reporter P14+ T cells.

**Supplementary fig. 5.**
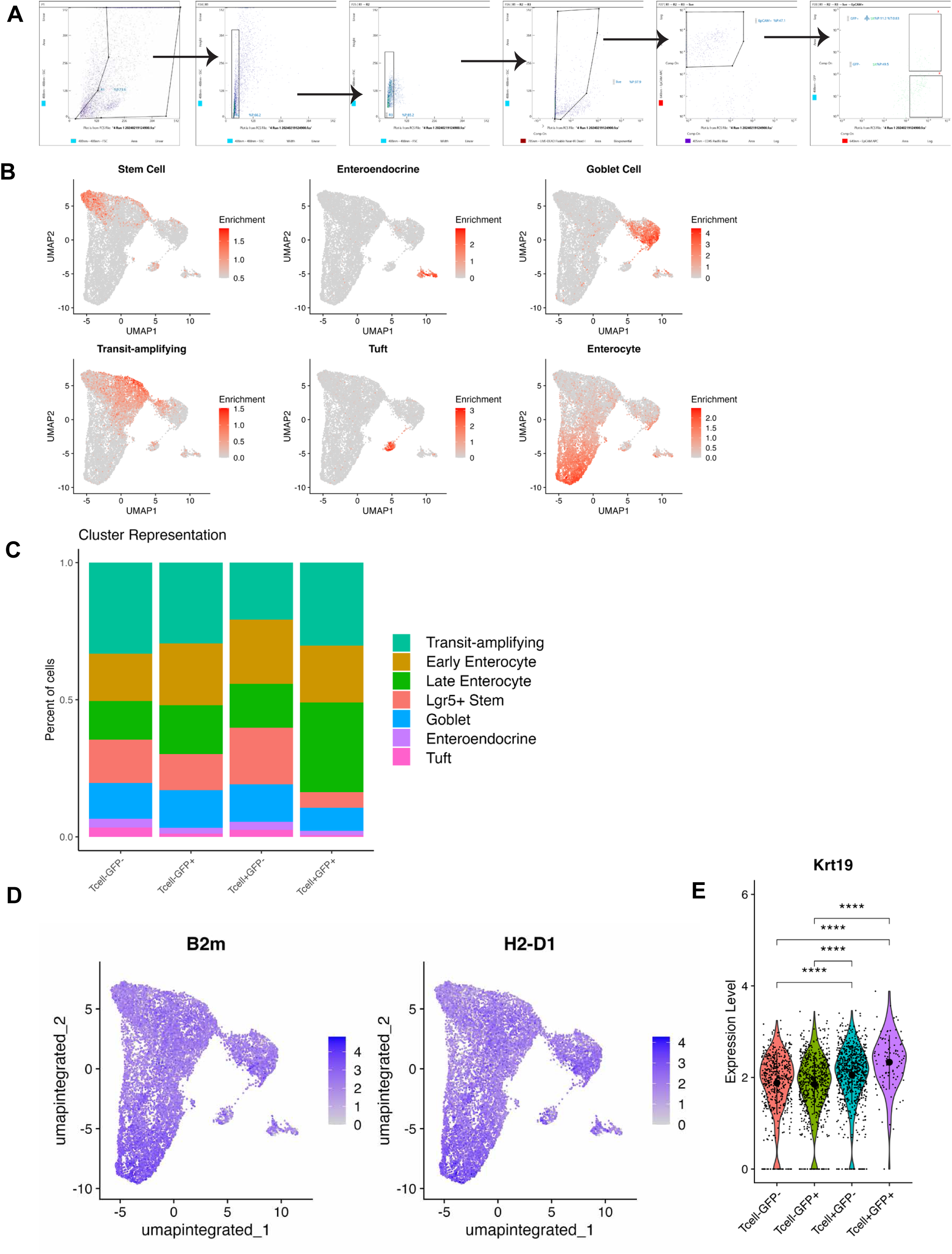
**a**,Gating strategy for the sorting of NINJA+ and NINJA-epithelial cells for scRNAseq. **b**, UMAP reduction with gene module enrichment, cell-type gene modules used for cluster definitions. **c**, Percent of cells, within each experimental group, in each cluster type, depicted as a stacked bar graph. **d**, UMAP reduction with B2M and H2-D1 expression. **e**, Violin plot of Krt19 transcript expression. A Kruskal-Wallis with Dunn’s multiple comparison test was performed to determine statistical significance. *P < 0.05; **P < 0.01; ***P < 0.001; ****P < 0.0001.

**Supplementary fig. 6.**
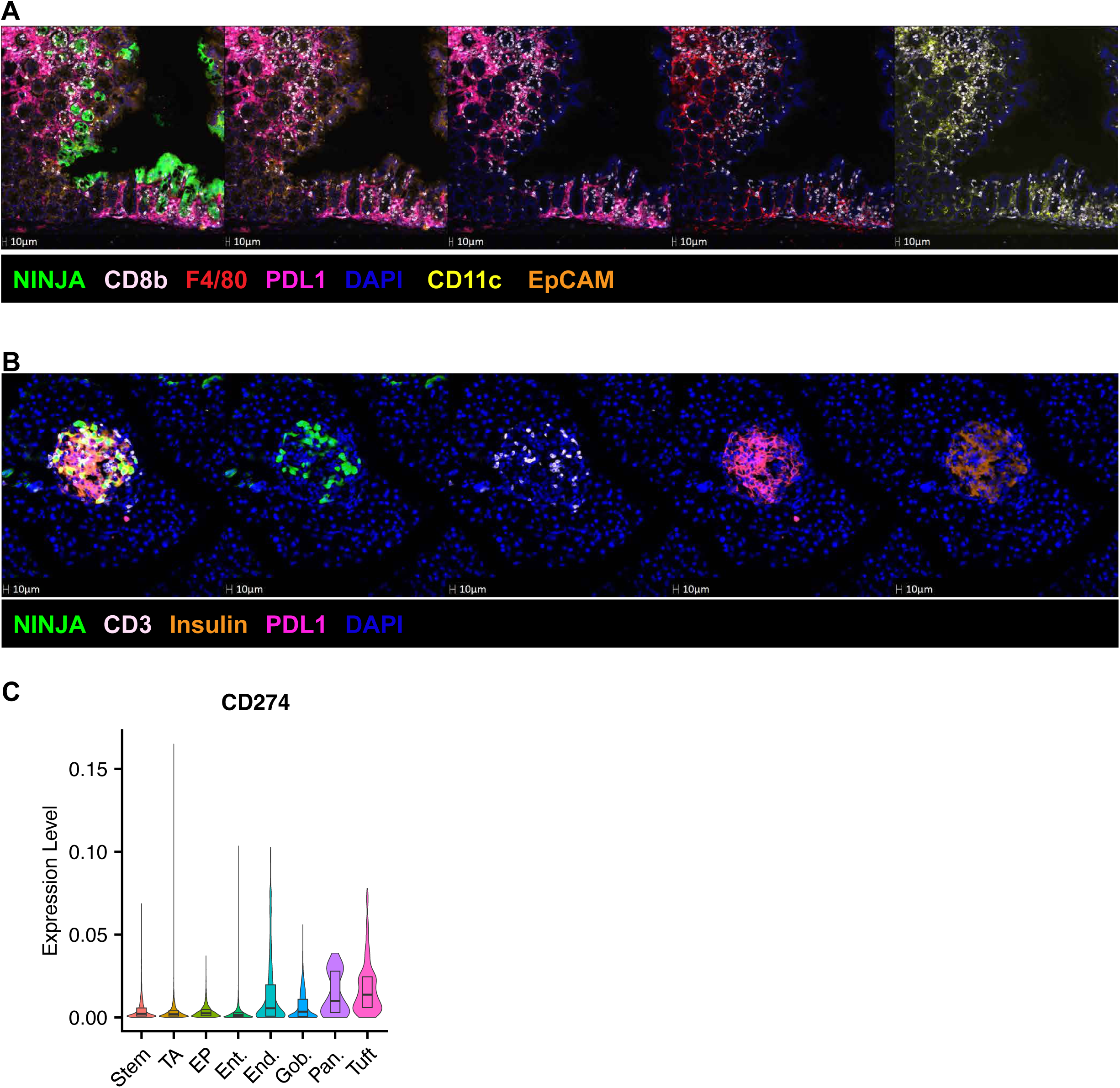
**a**, Zoomed out image of **Fig. 5a**, localization of PDL1, F4/80, and CD11c near NINJA+ epithelium. 20x CellDIVE immunofluorescence image. Green = NINJA and Nur77, white = CD8b, red = F4/80.1, pink = PDL1, yellow = CD11c, orange = EpCAM. **b**, representative 20x CellDIVE immunofluorescence image of T cell-infiltrated NINJA+ islets. Green = NINJA, white = CD3, pink = PDL1, orange = insulin, blue = DAPI. n = 15, accross 2 experiments **c**, Violin plot of PDL1 transcript expression in various C57BL/6 mouse small intestine epithelial cell populations.

**Supplementary fig. 7.**
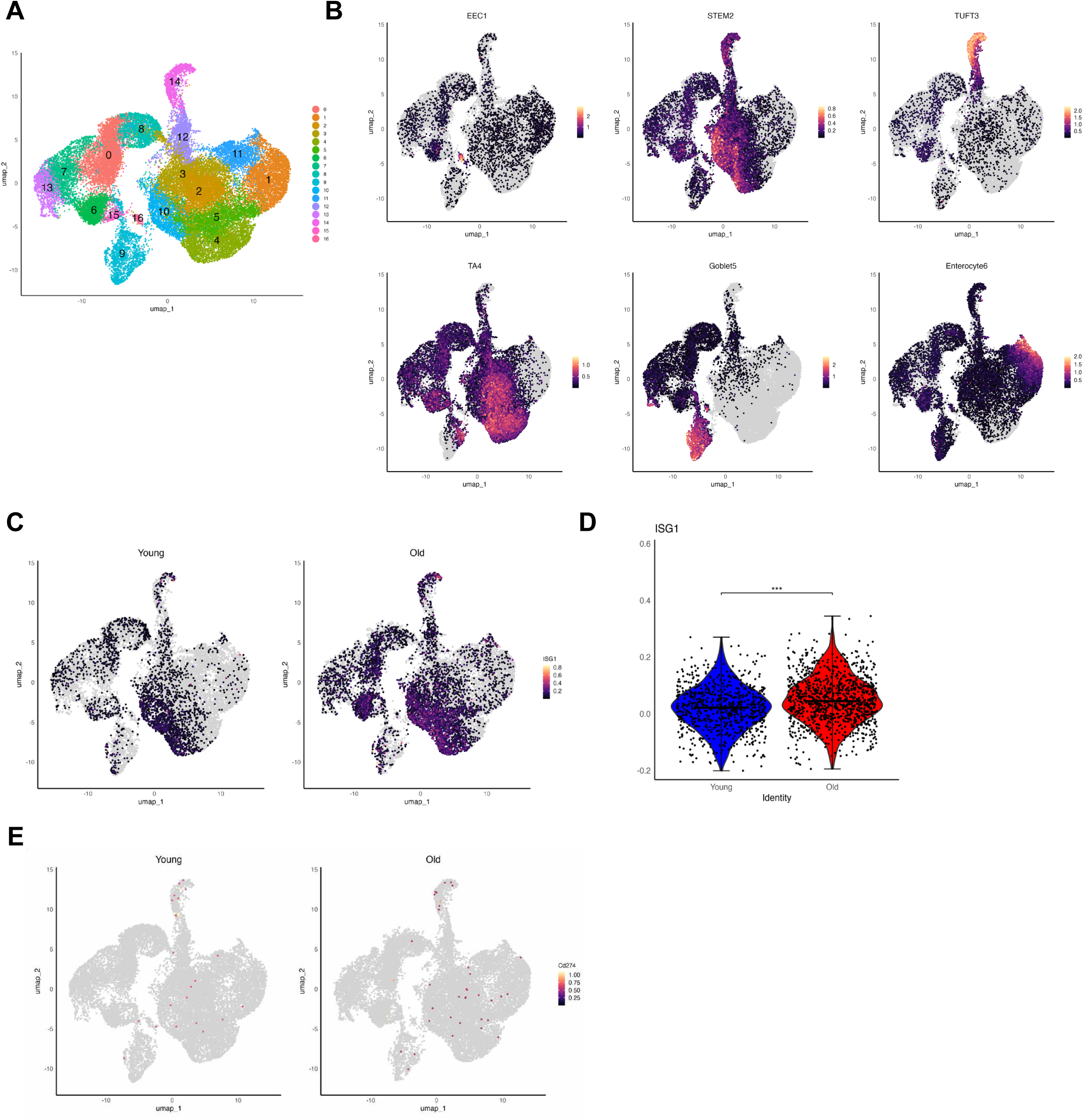
**a**,UMAP reduction with cluster definitions. **b**, UMAP reduction with expression of the gene modules used for cluster definitions. **c**, UMAP reduction of young and aged colon with ISG gene module expression. **d**, Violin plot of ISG gene module expression. Student’s t-test was performed to determine statistical significance. **e,** UMAP reduction of young and aged colon with CD274 transcript expression.

**Supplementary fig. 8.**
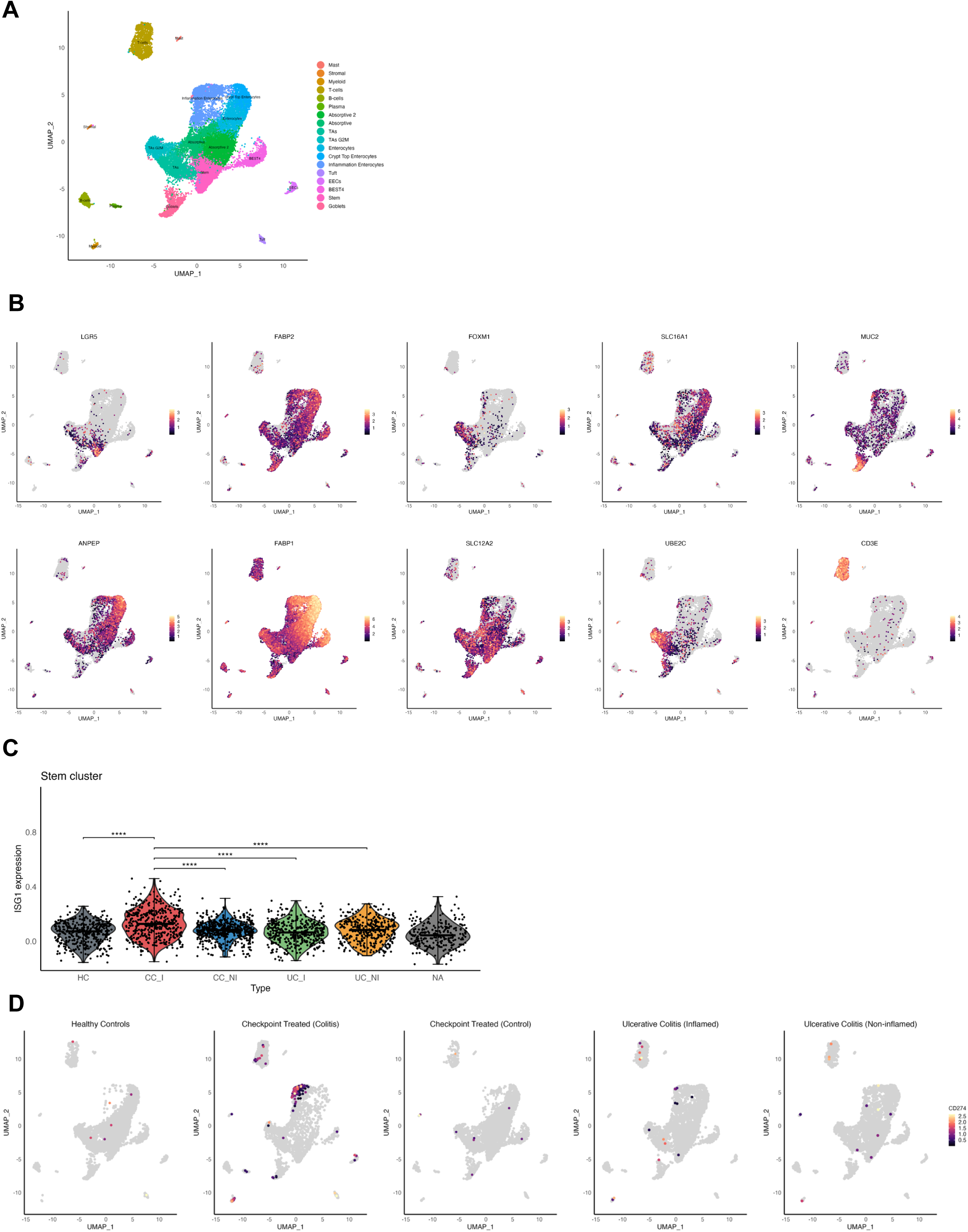
**a**, UMAP reduction with cluster definitions. **b**, UMAP reduction with expression of common cluster-specific genes. **c,** Violin plot of ISG gene module expression. A Kruskal-Wallis test with Dunn’s multiple comparison was performed to determine statistical significance. **d**, UMAP reduction with CD274 transcript expression in various conditions.

**Supplementary fig. 9.**
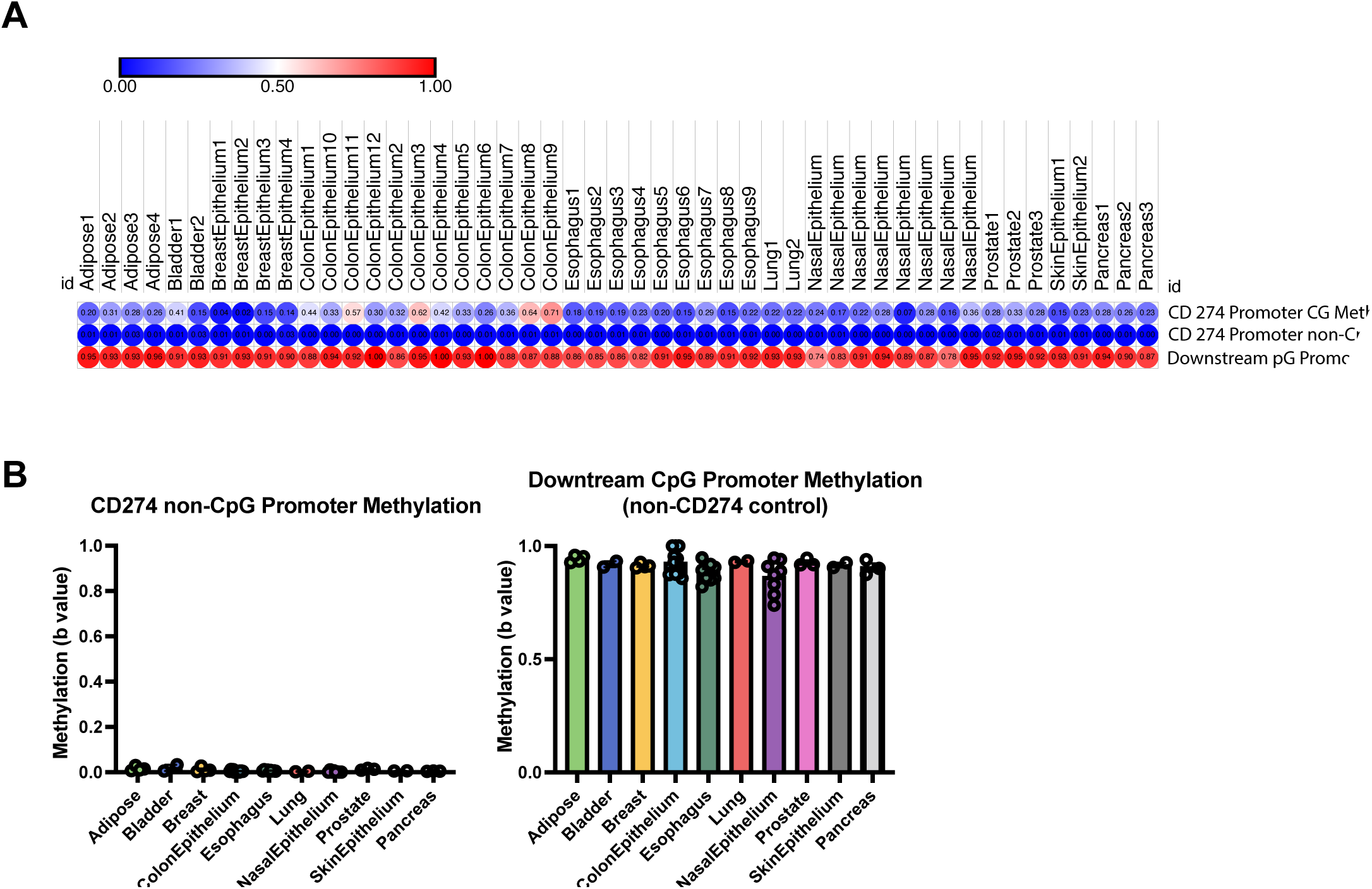
**a**,Bubble plot of PDL1 methylation in various tissues and locations. **b**, bar graph depicting non-CpG CD274 methylation and downstream CpG promoter methylation.

